# The conserved protein CBA1 is required for vitamin B_12_ uptake in different algal lineages

**DOI:** 10.1101/2023.03.24.534157

**Authors:** Andrew P. Sayer, Marcel Llavero-Pasquina, Katrin Geisler, Andre Holzer, Freddy Bunbury, Gonzalo I. Mendoza-Ochoa, Andrew D. Lawrence, Martin J. Warren, Payam Mehrshahi, Alison G. Smith

## Abstract

Microalgae play an essential role in global net primary productivity and global biogeochemical cycling, but despite their phototrophic lifestyle, over half of algal species depend on a supply of the corrinoid vitamin B_12_ (cobalamin) for growth. This essential organic micronutrient is produced only by a subset of prokaryotic organisms, which implies that for algal species to use this compound, they must first acquire it from external sources. Previous studies have identified protein components involved in vitamin B_12_ uptake in bacterial species and humans. However, little is known about how it is taken up in algae. Here, we demonstrate the essential role of a protein, CBA1 (for cobalamin acquisition protein 1), in B_12_ uptake in *Phaeodactylum tricornutum*, using CRISPR-Cas9 to generate targeted knockouts, and in *Chlamydomonas reinhardtii*, by insertional mutagenesis. In both cases, CBA1 knockout lines are no longer able to take up exogenous vitamin B_12_. Complementation of the *C. reinhardtii* mutants with the wildtype *CBA1* gene restores B_12_ uptake, and regulation of *CBA1* expression via a riboswitch element can be used to control the phenotype. When visualised by confocal microscopy, a YFP-fusion with *C. reinhardtii* CBA1 shows association with membranes. A bioinformatics analysis found that CBA1-like sequences are present in all the major eukaryotic phyla. Its presence is correlated with B_12_-dependent enzymes in many, although not all, taxa, suggesting CBA1 has a conserved role. Our results thus provide insight into the molecular basis of algal B_12_ acquisition, a process that likely underpins many interactions in aquatic microbial communities.

**One sentence summary:** Knockout mutants and physiological studies demonstrate that the CBA1 protein is essential for uptake of vitamin B_12_ in both *Chlamydomonas reinhardtii* and the unrelated *Phaeodactylum tricornutum*.

## INTRODUCTION

Microalgae are a diverse group of eukaryotic organisms that thrive in all aquatic environments. They form the basis of most aquatic food chains and are major contributors to global primary productivity, with marine microalgae responsible for an estimated 30% of total carbon fixation (Field et al., 1998). Understanding the drivers that support algal growth is thus of considerable ecological importance. Despite their photoautotrophic lifestyle, a widespread trait in algae is dependence on an external source of an organic micronutrient, vitamin B_12_ (cobalamin), a complex cobalt-containing corrinoid molecule. Approximately half of algal species surveyed across the eukaryotic tree of life require B_12_ for growth (Croft et al., 2005). However, the proportion of B_12_-dependent species differs between algal groups, from 30% (n=148) of Chlorophytes to 96% (n=27) of algal species that participate in harmful algal blooms (Tang et al., 2010). Within algal lineages, there is no evidence that any can produce B_12_ *de novo*, so this auxotrophy is not due to loss of one or more biosynthetic genes. Rather, the requirement for B_12_ stems from the fact that it is an essential cofactor for methionine synthase (METH), and species that can grow without supplementation have an alternative, B_12_-independent, isoform of this enzyme called METE (Croft et al., 2005; Helliwell et al., 2011). Many microalgae, including the green alga *Chlamydomonas reinhardtii* and the unrelated diatom *Phaeodactylum tricornutum*, encode both forms of methionine synthase and utilise METE in the absence of exogenous B_12_, but take up and utilise the compound if it becomes available (Helliwell et al., 2011). Under those conditions, the expression of *METE*, which has been found to have a lower catalytic rate than METH (Gonzalez et al. 1992), is repressed, and cells rely on METH activity.

The biosynthetic pathway for B_12_ is confined to prokaryotes (Warren et al., 2002) and indeed only a subset of bacteria encode the entire set of 20 or so enzymes required to synthesise corrinoids from the common tetrapyrrole precursor (Shelton et al., 2019), with many eubacterial species also reliant on an external source. In some cases, this is due to the loss of one or a few enzymes of the biosynthetic pathway, but in many bacteria the pathway is absent altogether and auxotrophy is the consequence of relying on one or more B_12_-dependent enzymes, such as METH. In microalgae, supplementation of cultures of *P. tricornutum* with B_12_ increases its growth rate subtly (Bertrand et al., 2012) and in *C. reinhardtii* use of METH confers thermal tolerance (Xie et al., 2013). More direct evidence for a selective advantage is demonstrated by the fact that an experimentally-evolved *metE* mutant of *C. reinhardtii* predominates in mixed populations with wild-type cells over tens of cell generations, as long as B_12_ is included in the medium (Helliwell et al., 2015). This is despite the fact that in the absence of B_12_, the *metE* mutant is non-viable within a few days (Bunbury et al., 2020).

The minimum levels of B_12_ in the medium needed to support growth of laboratory cultures of algal B_12_-auxotrophs are in the range of 10-50 pM (Croft et al., 2005), whereas B_12_ concentrations have been reported to be just 5-13 pM in freshwater systems (Ohwada, 1973). A similar value of 6.2 pM is the average value in most marine environments, although up to 87 pM could be detected in some coastal waters (Sañudo-Wilhelmy et al., 2014), which may be linked to the higher cobalt concentrations measured there (Panzeca et al., 2009). Given the limiting levels of B_12_ in the environment, its relatively short half-life (in the order of days) in surface water (Carlucci et al., 2007; Sañudo-Wilhelmy et al., 2014), and that as a large polar molecule it is unlikely to simply diffuse across cellular membranes, it is clear that algae must have an efficient means to take up B_12_. In bacteria, the molecular mechanisms for B_12_ uptake have been extensively characterised. The B_12_ transport and utilisation (*btu*) operon is perhaps the best known (Kadner, 1990), comprising BtuB, a TonB-dependent transporter in the outer membrane, a B_12_-binding protein, BtuF, located in the periplasm, and BtuC and BtuD, components of an ATP-binding cassette (ABC) transporter that sits in the inner membrane (Borths et al., 2002). In mammals, dietary B_12_ is bound to intrinsic factor in the ileum and taken up from the gut via receptor-mediated endocytosis (Nielsen et al., 2012). It is then transported between and within cells via multiple B_12_ transport proteins (Banerjee et al., 2021; Choi and Ford, 2021). These include LMBD1/ABCD4, the latter being an integral membrane ABC transporter in the lysosomal membrane of gut epithelial cells, which facilitates delivery of B_12_ into the cytosol, and MRP1 (or ABCC1), another ABC transporter that has sequence similarity to BtuCD and is involved in export of free B_12_ into the plasma where it binds to the main B_12_ transport protein, transcobalamin (Beedholm-Ebsen et al., 2010). Mice *mrp1* mutants were still able to transport a small amount of cobalamin out of cells, indicating redundant mechanisms for this function that have not yet been identified. Cobalamin circulating in the plasma bound to transcobalamin can then be taken up by other cells via receptor-mediated endocytosis (Nielsen et al., 2012).

In contrast to these well-studied processes in bacteria and mammals, the understanding of B_12_ acquisition in microalgae is more limited. A survey of microalgal species, including marine and freshwater taxa and those that require B_12_ (for example *Euglena gracilis*, *Thalassiosira pseudonana*) and non-requirers (such as *P. tricornutum*, *Dunaliella primolecta*), found that many released a ‘B_12_-binder’ into the medium, likely a protein, that appeared to sequester B_12_ from solution and thereby inhibited growth of B_12_-dependent algae (Pintner and Altmeyer, 1979). Its role was unknown, but it was postulated that it might be involved in competition for resources between microalgal species in the environment. Subsequently, a protein was purified from the medium of cultures of *T. pseudonana* with a high affinity binding constant of 2 pM for B_12_ (Sahni et al., 2001). In its native state it was an oligomer of >400 kDa, with subunits of ~80 kDa and the amino acid profile was determined, but it was not possible to obtain sufficient amounts to characterise further. A different approach was taken by Bertrand et al. (2012), who conducted a transcriptomics and proteomics study of *P. tricornutum* and *T. pseudonana* grown under low or sufficient B_12_ conditions. This led to the identification of a gene highly upregulated at the transcript and protein level in the absence of B_12_. Overexpression of this protein in *P. tricornutum* resulted in an increase in the rate of B_12_ uptake, and the protein was named <u>C</u>o<u>B</u>alamin <u>A</u>cquisition protein <u>1</u> (CBA1) although no direct role was established. In this study we have taken a mutagenesis approach to try to identify genes responsible for B_12_ uptake in both *P. tricornutum* and *C. reinhardtii*, including extending the work on CBA1. In addition, we have determined the extent to which candidate proteins are conserved throughout the algal lineages, making use of recent increases in algal sequencing data.

## RESULTS

### *P. tricornutum CBA1* knockout lines do not take up B_12_

Previous work showed that overexpression of CBA1 in *P. tricornutum* conferred enhanced B_12_ uptake rates (Bertrand et al., 2012) but the study did not demonstrate whether it was essential for this process. To address this question, *CBA1* knockout lines were generated in *P. tricornutum* strain 1055/1 (Table S1) by CRISPR-Cas9 editing, using a homologous recombination repair template that included a nourseothricin resistance (*NAT*) cassette (Figure 1a). CRISPR-Cas9 lines were cultured on selective media and screened for the absence of WT alleles at the *PtCBA1* locus (Phatr3_J48322) using PCR (Figure 1b). When the *PtCBA1* gene was amplified (top panel, Figure 1b) from ΔCBA1-1 with primers flanking the homologous recombination regions, two bands were detected; the larger of these corresponded to the WT amplicon, whilst the smaller band corresponded to a replacement of *CBA1* by *NAT,* suggesting that this strain is a mono-allelic knockout. For ΔCBA1-2, the *PtCBA1* gene primers amplified a single smaller product, suggesting that this was a bi-allelic knockout, whereas the *PtCBA1* ORF primers (bottom panel of Figure 1b) did not amplify anything, indicating a disruption specifically in this region. Similarly, no band was detected with primers that amplify across the 5’ end of the NAT knock-in (HR primers), which might indicate further disruptions upstream of the 5’HR region of ΔCBA1-2. Although a larger band than for WT was amplified in ΔCBA1-3 using the *PtCBA1* gene primers, those for the *PtCBA1* ORF amplified a smaller product; in both cases a single band was observed indicating a bi-allelic deletion at the sgRNA target sites.

**Figure 1.**
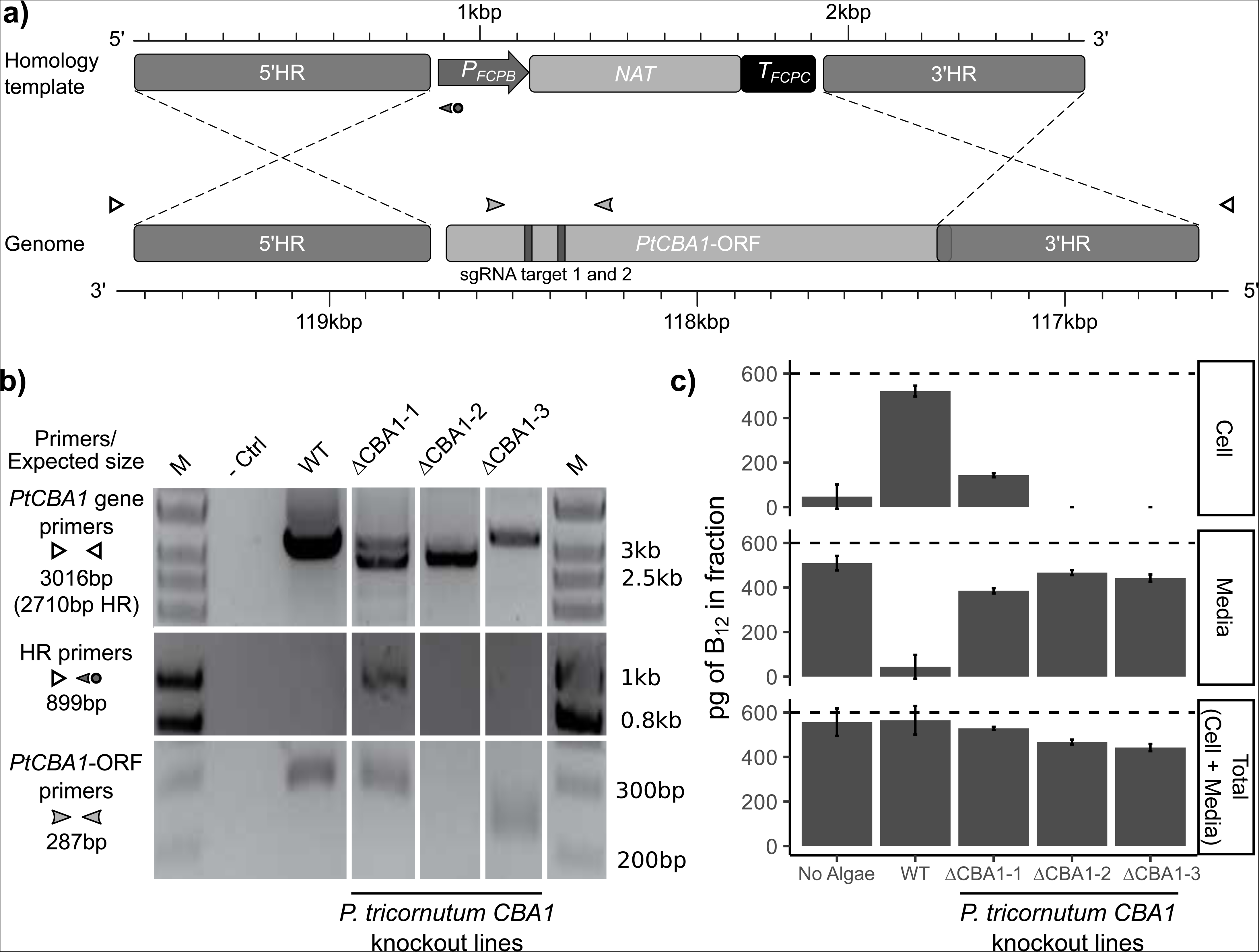
Disruption of *Phaeodactylum tricornutum CBA1* (*PtCBA1*) using CRISPR-Cas9 yielded lines with impaired B_12_ uptake. **a)** Schematic showing CRISPR-Cas9 sgRNA target sites and the homology repair template design used to generate mutant lines in *PtCBA1* (Phatr3_J48322). The homology repair template schematic is annotated with the 5’ homology region (HR) and 3’HR, the *FCPB* promoter, nourseothricin resistance gene (*NAT*) and *FCPC* terminator. The *PtCBA1* gene is annotated with the ORF, the 5’HR and 3’HR regions used in the homology template and the regions of the ORF targeted by sgRNA (vertical bars). Primer positions used for the analysis of putative mutant lines are shown with arrowheads. **b)** PCR of regions across and within wild-type (WT) and mutant *PtCBA1* in 3 independent CRISPR-Cas9 lines (ΔCBA1) showing indel mutations in the mutants. PCR products from different sets of primers indicated in panel a are shown. M = marker, - Ctrl = no DNA template. **c)** A B_12_ uptake assay was performed as described in Materials and Methods, to determine the amount of B_12_ in the media and the cells after 1h incubation of *P. tricornutum* cells in 600 pg B_12_. The ‘Total’ was inferred by the addition of the cell and media fractions. The dashed line indicates the amount of B_12_ added to the experiment. Standard deviation error bars are shown, n=4. Statistical analysis was performed on the media fraction, and Tukey’s test identified the following comparisons to be significantly different from one another: WT vs No Algae (p<1e^−12^); WT vs ΔCBA1-1 (p<1e^−10^); WT vs ΔCBA1-2 (p<1e^−12^); WT vs ΔCBA1-3 (p<1e^−11^); No Algae vs ΔCBA1-1 (p<1e^−03^); No Algae vs ΔCBA1-3 (p<0.05); and ΔCBA1-1 vs ΔCBA1-2 (p<1e^−02^).

To test whether the ΔCBA1 lines were affected in their ability to take up vitamin B_12_ we developed a standardised B_12_-uptake assay, detailed in Materials and Methods. In brief, algal cells were grown to the same growth stage and adjusted to the same cell density, then incubated in media containing a known amount of cyanocobalamin for one hour. Thereafter, cells were pelleted by centrifugation and the amount of B_12_ determined in the cell pellet and the media fraction using a *Salmonella typhimurium* bioassay (Bunbury et al., 2020). For each sample, the B_12_ measured in the cellular and media fractions were added to provide an estimated ‘Total’ and compared to the amount of B_12_ added initially (Figure 1c, dashed line), to determine the extent of recovery. For the WT strain, most of the added B_12_ was found in the cellular fraction. The mono-allelic knockout line ΔCBA1-1 consistently showed ~20-30% B_12_ uptake relative to the WT strain. This suggested that a single copy of *PtCBA1* is sufficient to confer B_12_ uptake in *P. tricornutum*, but not to the same extent as the WT strain. In contrast, for the two bi-allelic knockout lines (ΔCBA1-2 and ΔCBA1-3) no B_12_ was detected in the cellular fraction in any experiment, indicating that vitamin B_12_ uptake was fully impaired in the absence of a functional *PtCBA1* copy, at least at the limit of detection of the B_12_ bioassay (of the order of 10 pg). These results expand our understanding of *PtCBA1* by demonstrating that its presence is essential for B_12_ uptake and indicates that there is no functional redundancy to *PtCBA1*.

### Insertional mutagenesis identified the *C. reinhardtii* homologue of *CBA1*

Bertrand et al. (2012) reported that there were no detectable *CBA1* homologues in algal lineages outside the Stramenopiles, so to investigate B_12_ uptake in *C. reinhardtii*, we decided to take an insertional mutagenesis approach. We took advantage of the fact that B_12_ represses expression of the *METE* gene at the transcriptional level via the promoter (*P_METE_*), and that reporter genes driven by this genetic element respond similarly (Helliwell et al., 2014), to develop a highly sensitive screen for lines no longer able to respond to B_12_. We hypothesised that, since *P_METE_* is likely to respond specifically to intracellular B_12_, *P_METE_* would not be repressed in strains unable to take up B_12_ from the media, so the reporter would be expressed and functional. If the reporter were an antibiotic resistance gene, this would allow identification of B_12_ uptake mutants in a more high-throughput manner than the B_12_-uptake assay. The background strain for insertional mutagenesis was made by transforming *C. reinhardtii* strain UVM4 (Neupert et al., 2009) with plasmid pAS_R1 containing a paromomycin resistance gene (*aphVIII*) under control of *P_METE_* (Figure 2a, top construct). Lines of this strain were tested for their responsiveness to B_12_ and paromomycin. One line, UVM4-T12, showed the appropriate sensitivity with increasing repression of growth in paromomycin as B_12_ concentrations were increased, the effect being more marked at 15-20 µg·ml^−1^ paromomycin than at 5-10 µg·ml^−1^ (Figure 2b). This line thus allowed for an easily quantifiable growth phenotype that was proportionally related to B_12_ concentration.

**Figure 2.**
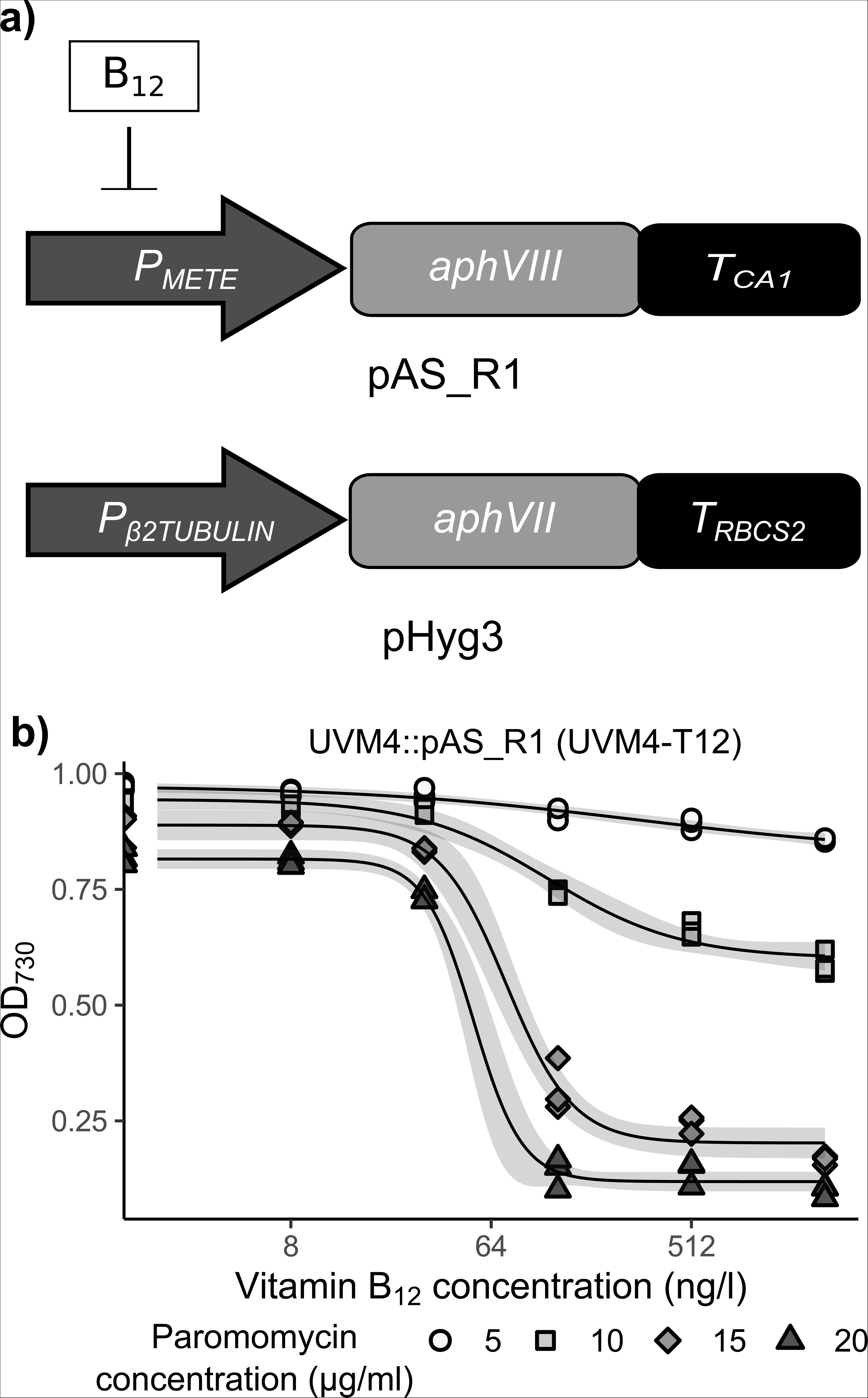
Generation and use of *C. reinhardtii* reporter strain UVM4-T12 for insertional mutagenesis. **a)** Schematic of the constructs used for insertional mutagenesis of *C. reinhardtii*. The pAS_R1 construct was designed to control expression of the paromomycin resistance gene (*aphVIII*) via B_12_ mediated repression of the *METE* promoter (*P_METE_*). The pHyg3 construct encoded a constitutively expressed hygromycin resistance gene (*aphVII*), to be used for insertional mutagenesis. **b)** Growth of *C. reinhardtii* B_12_ reporter strain UVM4-T12 bearing pAS_R1 plasmid, in response to vitamin B_12_ and paromomycin concentration in the media according to the algal dose-response assay. The predicted dose-response model is shown in black, with 95% confidence intervals in grey.

Insertional mutagenesis was carried out by transforming UVM4-T12 with a plasmid (pHyg3) containing a hygromycin resistance gene (*aphVII)* under the control of the constitutively expressed β2-tubulin promoter (Figure 2a, bottom construct), generating a population of UVM4-T12::pHyg3 lines with the cassette randomly inserted into the nuclear genome. By plating the products of the transformation on solid TAP media supplemented with a range of paromomycin, hygromycin and vitamin B_12_ concentrations (see Methods), 7 colonies were obtained. This was estimated to be from approximately 5000 primary transformants, determined by plating the same volume on TAP plates with the antibiotics but without B_12_. These 7 putative insertional mutant (IM) lines were then assessed for their ability to take up B_12_ using the B_12_ uptake assay. For UVM4, UVM4-T12 and insertional lines from the plate without B_12_ (labelled Control 1-3), similar amounts of B_12_ were recovered from the cellular and media fractions (Figure S1). This was also the case for 6 of the IM lines, suggesting that they could still take up B_12_ and were likely false positives of the initial screen. However, no B_12_ could be detected in the cellular fraction of UVM4-T12::pHyg3 #IM4 (hereafter referred to as IM4), indicating that this mutant line did not take up B_12_.

To obtain independent corroboration that IM4 was impaired in B_12_ uptake, cells of this mutagenized line were incubated with a fluorescently-labelled B_12_ derivative, B_12_-BODIPY (Lawrence et al., 2018), and then imaged using confocal microscopy. *C. reinhardtii* cells were incubated in TAP medium without B_12_-BODIPY or with 1 µM B_12_-BODIPY for 1 hour, washed with fresh media and subsequently imaged. There was no signal detected in the channel used for B_12_-BODIPY (589 nm excitation; 607-620 nm detection) in samples without B_12_-BODIPY added (Figure S2, top two rows), indicating that the imaging protocol was specific to this compound. When B_12_-BODIPY was added, UVM4-T12 showed the B_12_-BODIPY signal located within the algal cell (Figure S2, third row), indicating that this signal could be effectively detected by the imaging protocol and that B_12_-BODIPY was being transported into the cells. In contrast, there was no B_12_-BODIPY signal in IM4 cells, supporting the hypothesis that B_12_ uptake was impaired in this mutant (Figure S2, bottom row). In addition, the response of the *METE* gene to B_12_ in IM4 was assessed by RT-qPCR. UVM4 and IM4 cultures were grown in media with or without addition of B_12_ for 4 days in continuous light, after which the cultures were harvested for RNA extraction and cDNA synthesis. As expected, *METE* was repressed in UVM4 in the presence of B_12_ compared to no supplementation (Figure 3a), whereas IM4 showed similar *METE* expression in both conditions. This provided further support for disrupted B_12_ uptake in this line.

**Figure 3.**
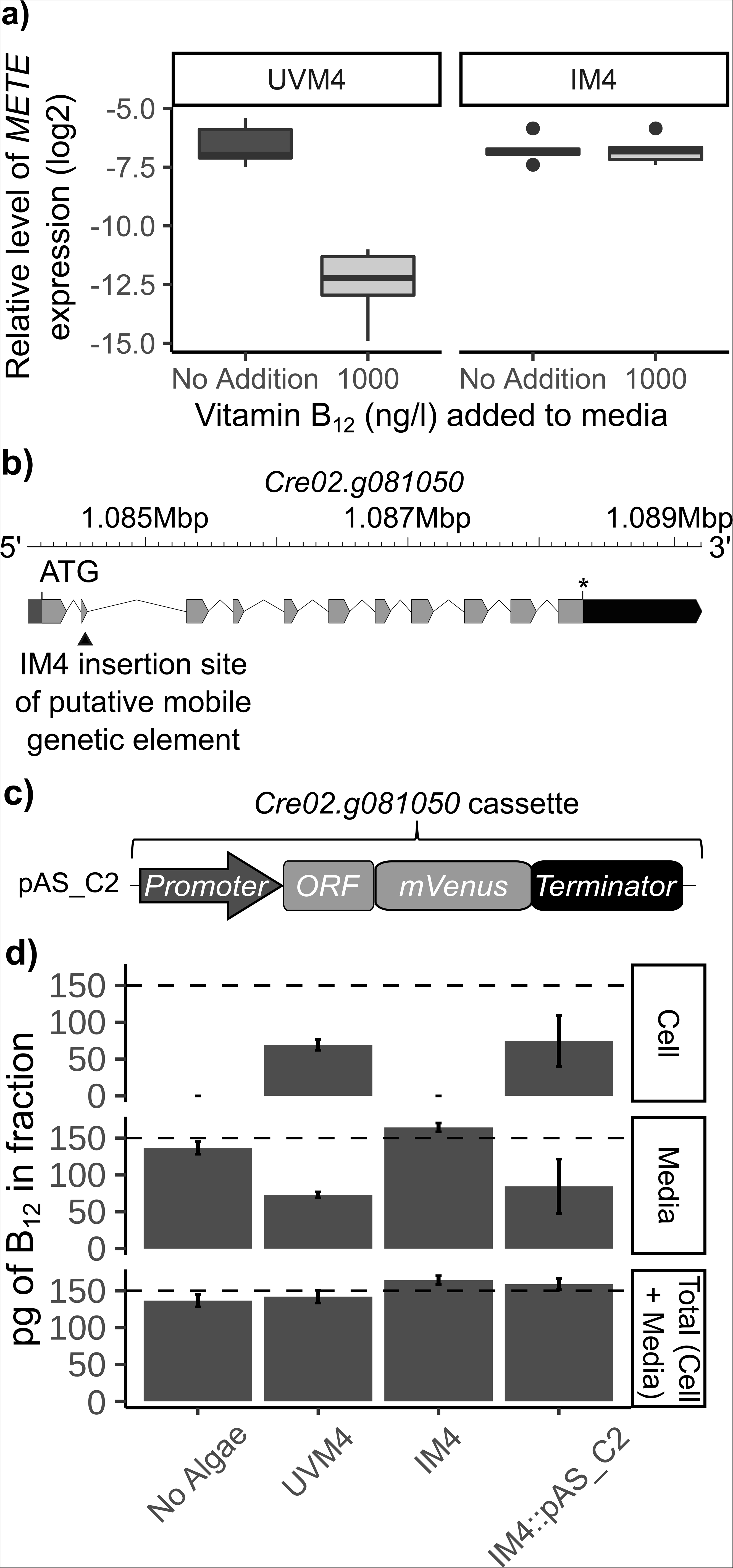
*C. reinhardtii* insertional mutant IM4 is defective in B_12_ response and uptake, and can be functionally complemented with *CrCBA1.* **a)** Effect of vitamin B_12_ on *METE* gene expression in UVM4 and IM4, determined by RT-qPCR. UVM4 and IM4 were grown in TAP media with or without 1000 ng·l^−1^ vitamin B_12_ for 4 days at 25°C, 120 rpm and in continuous light (90 µE·m^−2^·s^−1^). Boxplots of the log_2_ transformed relative expression level of *METE* to the *RACK1* housekeeping gene are shown, n=6. Significant comparisons were identified using Tukey’s test: UVM4 + 1000 ng·l^−1^ vitamin B_12_ from UVM4 No Addition (p<1e^−08^), IM4 No Addition (p<1e^−08^) and IM4 + 1000 ng·l^−1^ vitamin B12 (p<1e^−07^). **b)** Schematic of the Cre02.g081050 gene showing the position of the insertion site (indicated with a black triangle) determined by whole genome sequencing (Figure S4). **c)** Schematic of the pAS_C2 construct designed to express *CrCBA1* fused to the fluorescent reporter mVenus. *CrCBA1*-*mVenus* was under the control of the *CrCBA1* promoter and terminator. pAS_C2 also contained the spectinomycin resistance gene *aadA,* driven by the *PSAD* promoter and *PSAD* terminator. **d)** B_12_-uptake assay with UVM4, IM4 and IM4::pAS_C2 (n =4 separate transformants with high mVenus expression). Dashed line indicates the amount of B_12_ added to the assay. Standard deviation error bars are shown. Statistical analysis was performed on the media fraction, and Tukey’s test identified the following comparisons to be significantly different from one another: No Algae vs UVM4 (p<1e^−05^); No Algae vs IM4 (p<0.05); No Algae vs IM4::pAS_C2 (p<1e^−03^); UVM4 vs 1.G2 (p<1e^−09^); and 1.G2 vs 1.G2::pAS_C2 (p<1e^−06^).

To identify the genomic location of the causal mutation in IM4, short-read whole genome sequencing was performed on DNA samples from UVM4, UVM4-T12 and IM4. The location of the pHyg3 cassette in IM4 was identified as described in Methods and found to have disrupted the *Cre12.g508644* locus (Figure S3a), an unannotated gene. To corroborate that disruption of the *Cre12.g508644* was responsible for the uptake-phenotype, two independent mutant lines of the gene (LMJ-119922 and LMJ-042227) were ordered from the Chlamydomonas library project (CLiP) collection (Li et al., 2016) and verified to be disrupted at this locus by PCR (Figure S3a). However, when these knockout lines were tested for the ability to take up B_12_ using the B_12_ uptake assay, they were both found to be able to do so to a similar extent as their parental strain, cw15 (Figure S3b). This suggested that *Cre12.g508644* did not encode a protein essential for B_12_ uptake.

We therefore examined the genome sequence data more closely to determine the genetic cause for the B_12_-uptake phenotype of IM4. We had identified putative homologues of human proteins involved in receptor-mediated endocytosis of B_12_, such as ABCD4, LMBD1 (Rutsch et al., 2009; Coelho et al., 2012) and MRP1 (Beedholm-Ebsen et al., 2010), in the *C. reinhardtii* genome by BLAST (data not shown). However, given the widespread percentage of SNPs in the IM4 genome compared to UVM4, it was not possible to identify any candidate causal mutations with confidence. Instead, manual inspection of the DNA sequencing reads mapped to the reference strain revealed one locus, *Cre02.g081050,* annotated as flagella-associated protein 24 (FAP24), where there was a unique discontinuity, suggesting that there was an insertion at exon 2 in IM4 (Figure 3b; Figure S4a). The sequence was bordered by a genome duplication of 8 bp (shown in blue in Figure S4a) and exhibited imperfect inverted repeats at the terminal regions (TIRs), indicative of a transposable element. Reads could not be assembled across the discontinuity to obtain the complete sequence of the insertion, but using the left and right junction sequences as queries, three regions encoding two very similar genes were identified (Figure S4b).

Remarkably, when the *Cre02.g081050* protein was used as a query in a BLAST search, one of the hits recovered was the *PtCBA1* protein (22.9% sequence identity), even though the reciprocal sequence search had not picked up the *C. reinhardtii* gene (Bertrand et al., 2012). The Phyre2 structural prediction server (Kelley et al., 2015) was used to model the 3D structures of PtCBA1 and the *C. reinhardtii* protein encoded by *Cre02.g081050* (Figure S5). The modelled proteins showed a high degree of structural similarity to one another (root mean squared deviation (RMSD) = 2.333), particularly with respect to the arrangement of alpha helices and lower cleft. Due to the sequence similarity and predicted structural similarity, these proteins appeared to be homologous to one another and Cre02.g081050 is hereafter referred to as CrCBA1.

To determine whether disruption of *CrCBA1* in IM4 was responsible for the impaired B_12_ uptake, we investigated whether it was possible to restore its ability to take up B_12_ by transforming IM4 with the wild-type *CrCBA1*. Construct pAS_C2 was designed with the *CrCBA1* promoter, *CrCBA1* open reading frame (ORF) and terminator and included a 3’ mVenus tag attached by a poly-glycine linker (Figure 3c). IM4 was transformed with pAS_C2, and resulting lines were tested for the ability to take up B_12_ using the B_12_ uptake assay. As observed previously, UVM4 was able to take up B_12_ whilst IM4 was unable to do so (Figure 3d). The CBA1 complementation line IM4::pAS_C2 showed B_12_ in the cellular fraction at similar levels as in UVM4, thereby indicating that the mutant phenotype had been complemented.

### *CrCBA1* CLiP mutant is unable to take up B_12_ and is complemented by the WT *CrCBA1* gene

Given the many genetic changes in line IM4 compared to the parental UVM4-T12 strain caused by the mutagenesis, it was essential to have independent corroboration that mutation of *CrCBA1* caused the inability to take up B_12_. Accordingly, we obtained two further CLiP mutants (LMJ-135929 and LMJ-040682) with disruptions in intron 2 and introns 6/7 respectively of *CrCBA1* (Figure S6a) and assessed them for their ability to take up B_12_ (Figure S6b). No B_12_ was detected in cells of LMJ-040682, indicating complete inhibition of B_12_ uptake. Although LMJ-135929 cells accumulated some B_12_, this was less than half the amount of its parent strain cw15, suggesting partial impairment in uptake, similar to the phenotype of the monoallelic *PtCBA1* knockout line (Figure 1c). However, heterozygosity cannot be the explanation for *C. reinhardtii,* which is haploid, and instead indicates that LMJ-135929 was likely to have just partial knockdown of the gene, probably because the insertion is in an intron.

Nonetheless, to provide further confirmation that mutations in *CrCBA1* were responsible for the observed impaired B_12_ uptake, we again tested whether the phenotype could be complemented with the wild-type *CrCBA1* gene using both plasmid pAS_C2 (Figure 3b) and an additional construct pAS_C3 (Figure 4a), in which expression of *CrCBA1* can be controlled by a thiamine pyrophosphate (TPP) repressible riboswitch, RS*_THI4_4N_* (Mehrshahi et al., 2020). In the absence of thiamine supplementation of the cultures, the riboswitch is not active and the gene containing it is transcribed and translated as normal; with thiamine addition, alternative splice sites are utilised, leading to inclusion of an upstream ORF containing a stop codon in the mRNA, preventing translation from the downstream start codon. LMJ-040682 was transformed with both pAS_C2 and pAS_C3, and representative transformant lines selected via antibiotic resistance were obtained. These, together with their parental strains were grown in the presence or absence of 10 µM thiamine for 5 days, and then used in the B_12_ uptake assay. Transformants of both LMJ-040682::pAS_C2 and LMJ-040682::pAS_C3 were found to take up B_12_ to a similar extent as their parental strain cw15 when grown in the absence of thiamine (Figure 4b). However, when 10 µM thiamine was included in the culture medium, LMJ-040682::pAS_C3 showed virtually no B_12_ uptake. This riboswitch-mediated conditional complementation of the phenotype in LMJ-040682::pAS_C3 demonstrated conclusively that B_12_ uptake in *C. reinhardtii* is dependent on the presence of CrCBA1.

**Figure 4.**
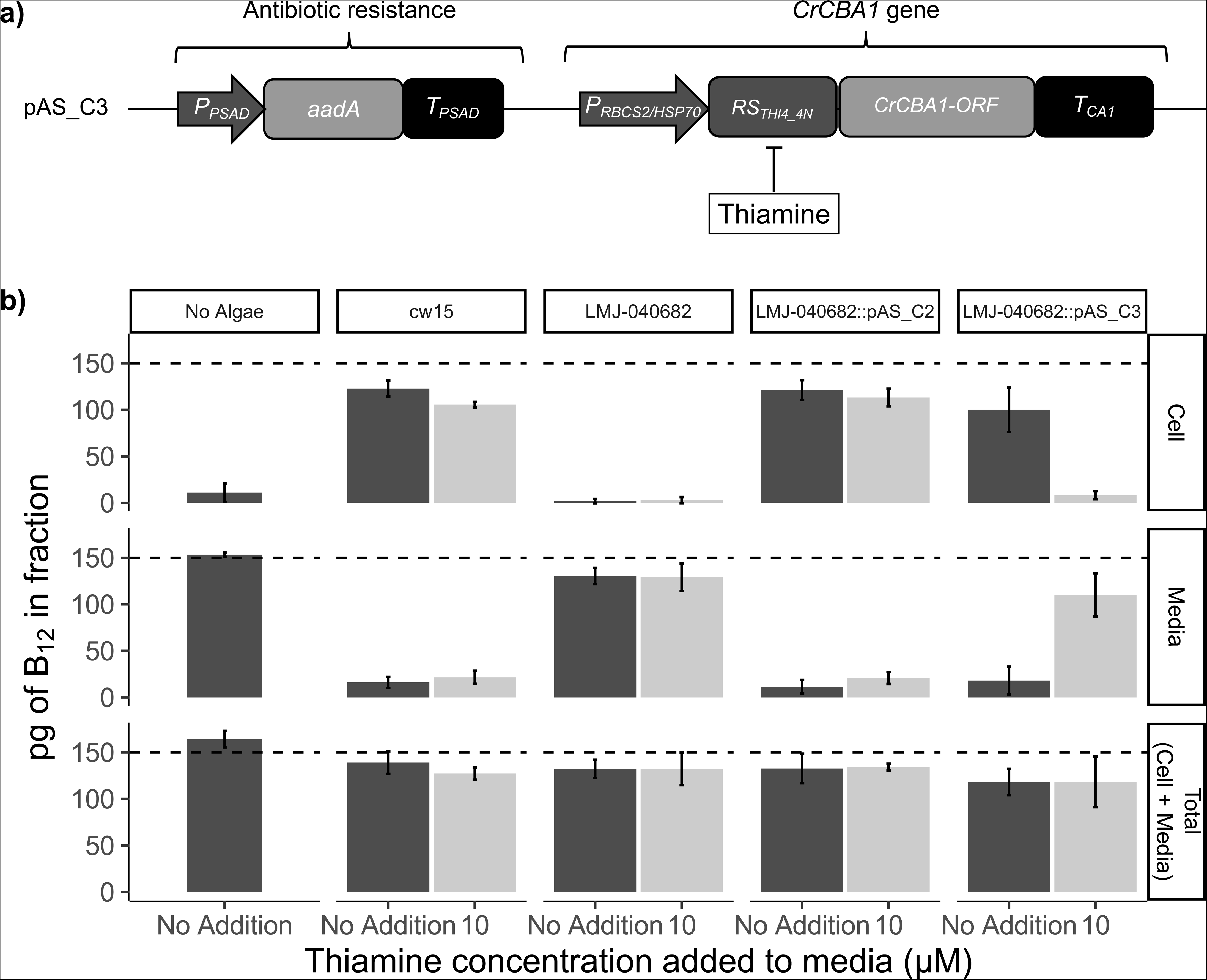
CLiP mutants in CrCBA1 are impaired in their ability to take up B_12_. **a)** Schematic of the pAS_C3 construct designed to express *CrCBA1* in a controllable manner using a thiamine repressible riboswitch (RS*_THI4_4N_*) to allow repression of *CrCBA1* through the addition of thiamine (Mehrshahi et al., 2020). **b)** B_12_-uptake assay with cw15, LMJ-040682 and mean of 3 independent transformants of LMJ-040682::pAS_C2 and LMJ-040682::pAS_C3. The growth conditions were modified compared to previous assays: lines were grown with or without 10 µM thiamine supplementation for 5 days in a 16/8 light/dark cycle, and 8 hours after the dark to light transition the cultures were used for the algal B_12_-uptake assay. The dashed line indicates the amount of B_12_ added to the sample. Standard deviation error bars are shown. Statistical analysis was performed on the media fraction. Tukey’s test identified the following algal strains to be significantly different from one another in media without thiamine (not reporting comparisons against the No Algae control condition): cw15 vs LMJ-040682 (p<1e^−10^); LMJ-040682 vs LMJ-040682::pAS_C2 (p<1e^−09^); and LMJ-040682 vs LMJ-040682::pAS_C3 (p<1e^−09^). Additionally, Tukey’s test found the following strain to show a significant difference due to thiamine addition: LMJ-040682::pAS_C3 (p<1e^−07^).

### CrCBA1 shows an association with membranes and is highly upregulated under B_12_-deprivation

To investigate the subcellular location of CrCBA1, we used several bioinformatic targeting prediction tools. CrCBA1 is annotated as a flagella-associated protein in the Phytozome v5.6 *C. reinhardtii* annotation. However, both DeepLoc (Almagro Armenteros et al., 2017) and SignalP (Almagro Armenteros et al., 2019) indicated a hydrophobic sequence with the characteristics of a signal peptide at the N-terminus of CrCBA1 and predicted it would be targeted to the endoplasmic reticulum (ER). Additionally, it was predicted to contain a transmembrane helix at its C-terminus by InterPro (Mitchell et al., 2019).

We next investigated the subcellular location of CrCBA1 *in vivo* by imaging two lines of LMJ-040682::pAS_C2, where the CBA1 is tagged with mVenus, with confocal microscopy. No mVenus was detected in the parental LMJ-040682 cells, whereas a clear fluorescent signal was observed in LMJ-040682::pAS_C2 #A10 and LMJ-040682::pAS_C2 #D10 (Figure 5). In these complemented lines, the mVenus signal was absent from the chloroplast, nucleus and flagella, but instead could be seen within the cell localising both to the plasma membrane and to regions that may be endomembranes such as the ER. This is consistent with findings from *P. tricornutum* showing a similar distribution (Bertrand et al., 2012). Together these data indicate that CBA1 is likely to be associated with membranes, and therefore, may have a conserved role in the B_12_ uptake process.

**Figure 5.**
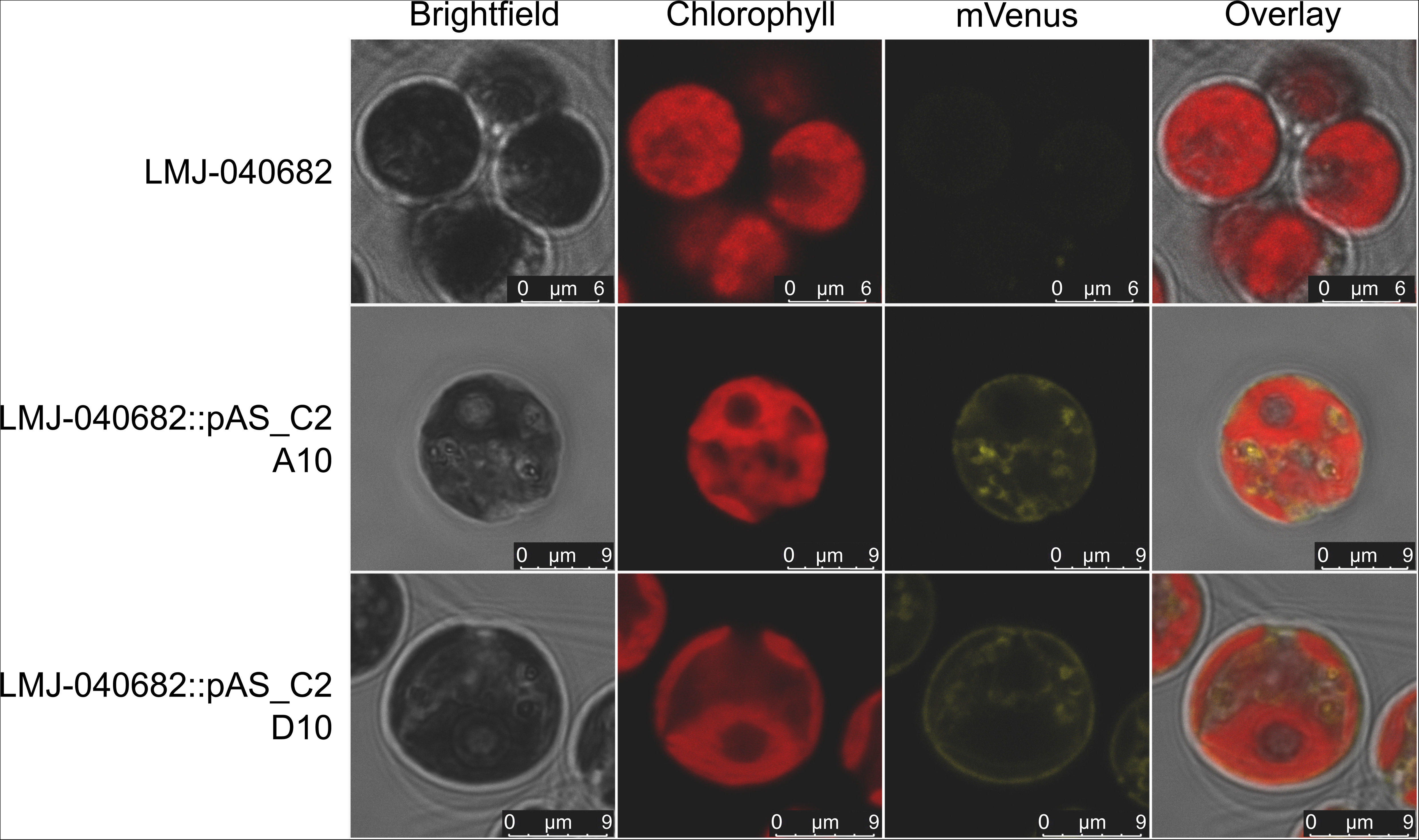
Confocal microscopy of complemented *C. reinhardtii CrCBA1* knockout lines showing an association between CrCBA1 and membranes. LMJ-040682 and LMJ-040682::pAS_C2 A10 and D10 lines were imaged according to the protocol outlined in the materials and methods. Channels shown (left to right) are brightfield, chlorophyll, mVenus and an overlay. Microscope settings are described in Methods.

Further evidence for the role of CBA1 in B_12_ uptake was obtained by taking advantage of a B_12_-dependent mutant of *C. reinhardtii*, metE7 (Helliwell et al., 2015; Bunbury et al., 2020). We tested the effect of B_12_-deprivation over time on the expression of the *CrCBA1* gene by RT-qPCR in the mutant and determined the rate of B_12_ uptake over a similar period. Within 6h of B_12_ removal, there was a ~250-fold induction of the *CrCBA1* transcript, followed by a slow decline over the next 60h (Figure 6a). After resupply of B_12_ there was then a rapid ~100-fold decline within 8h. The B_12_ uptake capacity of metE7 followed a similar profile, increasing 3-fold over the first 12 hours of B_12_ depletion, from ~6.5 x 10^5^ molecules B_12_/cell/hour to 1.86 x 10^6^ molecules B_12_/cell/hour (Figure 6b), then declining slowly. This induction profile is characteristic of a nutrient-starvation response shown by many transporters, including in *C. reinhardtii* those for Fe (Allen et al., 2007), and for *CBA1* in the B_12_-dependent diatom, *Thalassiosira pseudonana* (Bertrand et al., 2012).

**Figure 6.**
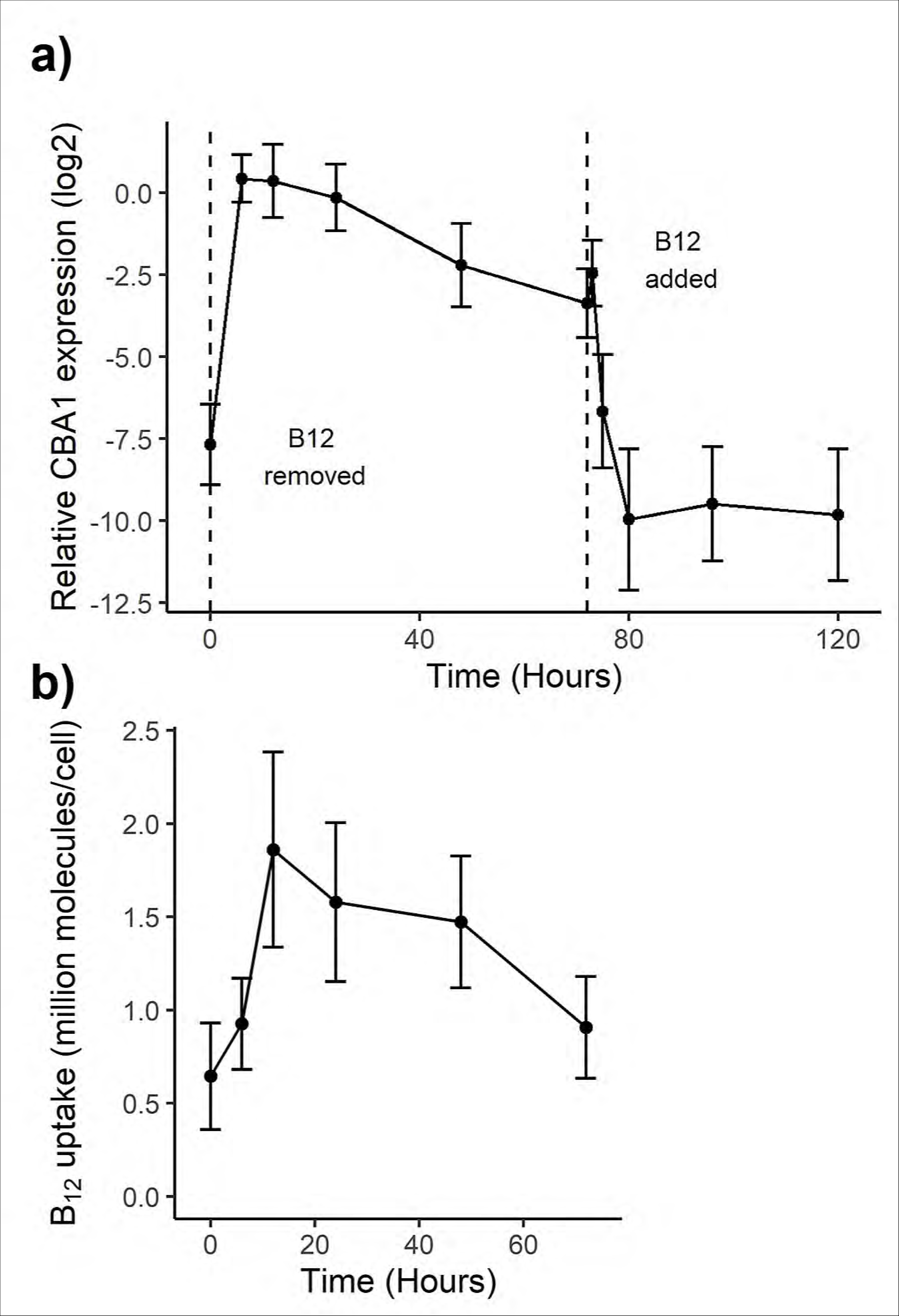
CBA1 expression and B_12_ uptake capacity in a B_12_-dependent mutant of C. reinhardtii (metE7) during B_12_ starvation and add-back. a) CBA1 expression measured by RT-qPCR and expressed relative to the housekeeping gene RACK1 using the 2^(ΔCt) method. Vertical dashed lines denote when B_12_ was removed and added. b) B_12_ uptake capacity of starved metE7 cells (expressed as 10^6^ molecules of B_12_ per cell over 1h) at the same 6 time points during B_12_ starvation; it was not possible to perform the uptake assay on cells to which B_12_ had already been added. Cell density measurements were performed by counting plated cells in dilution series, and so included non-viable cells. For CBA1 expression and B_12_ uptake, 3 and 6 biological replicates were used, respectively, with points representing means, and error bars representing standard deviations.

### Widespread distribution of CBA1 in algae

Having shown the importance of *PtCBA1* and *CrCBA1* for B_12_ uptake in their respective species, we re-examined how prevalent CBA1-like proteins are in Nature. Searches with BLASTP using *PtCBA1* resulted in no significant homologues in species outside the Stramenopiles (Bertrand et al., 2012). Instead, we created a hidden Markov model (HMM), using the *C. reinhardtii* CBA1 amino acid sequence and CBA1 sequences from *P. tricornutum*, *T. pseudonana*, *Fragilariopsis cylindrus*, *Aureococcus anophagefferens* and *Ectocarpus siliculosus* (Bertrand et al., 2012), to identify more accurately CBA1-like proteins in other organisms. The EukProt database of curated eukaryotic genomes (Richter et al. 2022) includes representatives from the Archaeplastida (designated by EukProt as Chloroplastida), which encompass green algae, red algae, glaucophytes and all land plants, as well as phyla that include algae with complex plastids, namely Stramenopiles (which include diatoms), Alveolata (including dinoflagellates), Rhizaria and Haptophyta, and the animals (both Metazoa and basal Choanoflagellates), the fungi and Amoebozoa. This database was queried with the CBA1 HMM model, using a cutoff e-value of 1e-20, and 277 hits were obtained (Figure S7; Supplementary Table S3). No candidates were found in the Metazoa, but CBA1 homologues were identified in all other phyla, including all photosynthetic groups, fungi and amoebozoa and in choanoflagellates, unicellular and colonial flagellated organisms considered to be the closest living relatives of animals (King et al., 2008).

Given that higher plants have no B_12_-dependent enzymes, the presence of a putative B_12_-binding protein in several angiosperms, both monocot and dicot, and the gymnosperm *Ginkgo biloba*, was somewhat surprising. To address this conundrum, we investigated to what extent CBA1 was associated with vitamin B_12_ dependence by determining the distribution of the different isoforms of methionine synthase, METH and METE. Using the same HMM approach as before, the protein sequences were searched against the EukProt database and the combination of presence and absence of CBA1, METH and METE across eukaryotic species groups was compiled (Figure 7; Supplementary Table S4). What is immediately apparent is that the combination of the three proteins is quite different in the various lineages. In the major algal groups, the Chlorophyta and the SAR clade (Stramenopiles, Alveolata and Rhizaria), METH sequences were found in the majority of genomes analysed and their presence was correlated with CBA1. In the genomes of the Chlorophyta and the SAR clade that encoded METE only (7 taxa in total), CBA1 was absent in all but one, the diatom *Thalassionema nitzschiodes*. Equal numbers of Alveolata species encoded METH and CBA1, or METH only; interestingly, the latter were all non-photosynthetic lineages. Grouping the data from these 4 algal groups, a Chi Square test was significant for CBA1 and METH being more often both present or both absent (*X2 (1, N = 86) = 9.2, p = 0.00240).* The association could be due to linkage, although in neither *C. reinhardtii* nor *P. tricornutum* are the two genes on the same chromosome, making this unlikely. Alternatively, there is a fitness advantage in both genes being acquired or lost together.

**Figure 7.**
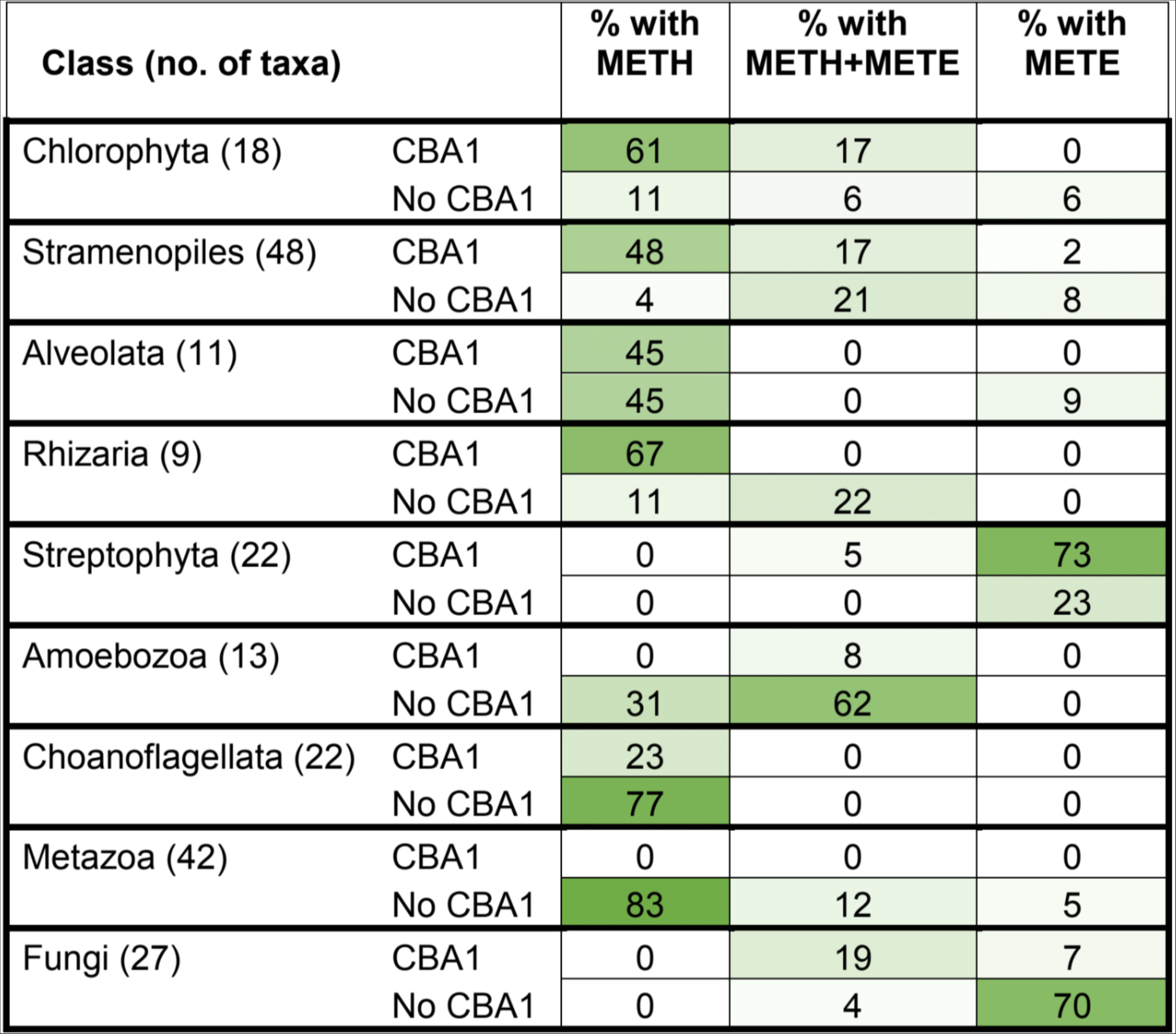
Distribution of CBA1 and methionine synthase sequences across Eukaryotic groups. The EukProt database (Richter et al., 2022) was searched for METE, METH and CBA1 queries, as described in the materials and methods. Organisms were only considered if they contained at least one valid methionine synthase hit (METE or METH) and their genomes were >70% complete, as measured by BUSCO (Manni et al., 2021). Eukaryotic classes were filtered for those with greater than 5 genomes and the numbers of taxa for each class are indicated in brackets. The different combinations of CBA1, METE and METH were calculated for each species (Supplementary Table 4) and summarised as a percentage of the total number of taxa in each class, with gradual shading to show the variation in distribution between the different classes.

Most fungal taxa lacked both METH and CBA1, but we found examples of 6 species that were predicted to be B_12_ users (METH present) and 5 of these were also predicted to contain CBA1-like sequences: *Allomyces macrogynus*, *Spizellomyces punctatus*, *Rhizophagus irregularis*, *Rhizopus delemar* and *Phycomyces blakesleeanus*. CBA1-like sequences were identified in the Opisthokonta and Amoebozoa, although were less prevalent, with ~23% of choanoflagellates and 8% of amoeboid species being like algae in having both METH and CBA1. CBA1 was entirely absent from the Metazoa. In contrast, in the Streptophyta, which include multicellular green algae and all land plants, the majority lack METH, but almost 80% of species were found to contain CBA1-like sequences. This implies that Streptophyta CBA1 sequences may have gained a different function, which would be consistent with the lack of B_12_-dependent metabolism in these organisms. In summary, these data suggest that CBA1 is associated with vitamin B_12_ use to different degrees in different eukaryotic groups, with there being a greater association in obligate and facultative B_12_ users than in those organisms that do not utilise B_12_.

The many putative CBA1 homologues in algal lineages and their strong association with B_12_ uptake provided an opportunity to identify conserved, and thus likely functionally important, residues. Accordingly, a multiple sequence alignment of proteins matching the CBA1 HMM query was generated (Figure S7). Highlighted in green in the similarity matrix at the top are nine conserved regions with several almost completely conserved residues; these are shown in more detail in Figure 8a for selected taxa representing different algal groups. Further insight came from inspection of the model of the 3D structure of CrCBA1 generated by the Phyre2 structural prediction server. The analysis showed that regions of CrCBA1 showed similarity to bacterial periplasmic binding proteins, including the B_12_-binding protein BtuF. A structure is available of *E. coli* BtuF in complex with B_12_ (Borths et al., 2002), so we compared this to the modelled CrCBA1 structure. Although there is little sequence similarity, alignment of the two structures resulted in an RMSD of 3.362 and enabled the relative position of B_12_ to be placed in the lower cleft of CrCBA1, shown in red in Figure 8b. Mapping of the highly conserved residues onto this structure found that many (P251, V253, W255, W394, F395 and E396) were in a cluster around the relative position of B_12_. Another cluster of highly conserved residues were located at the end of the upper alpha helix (P118, L136, F214, F215, N216 and E218). Both clusters represent promising mutational targets to investigate CrCBA1 function.

**Figure 8.**
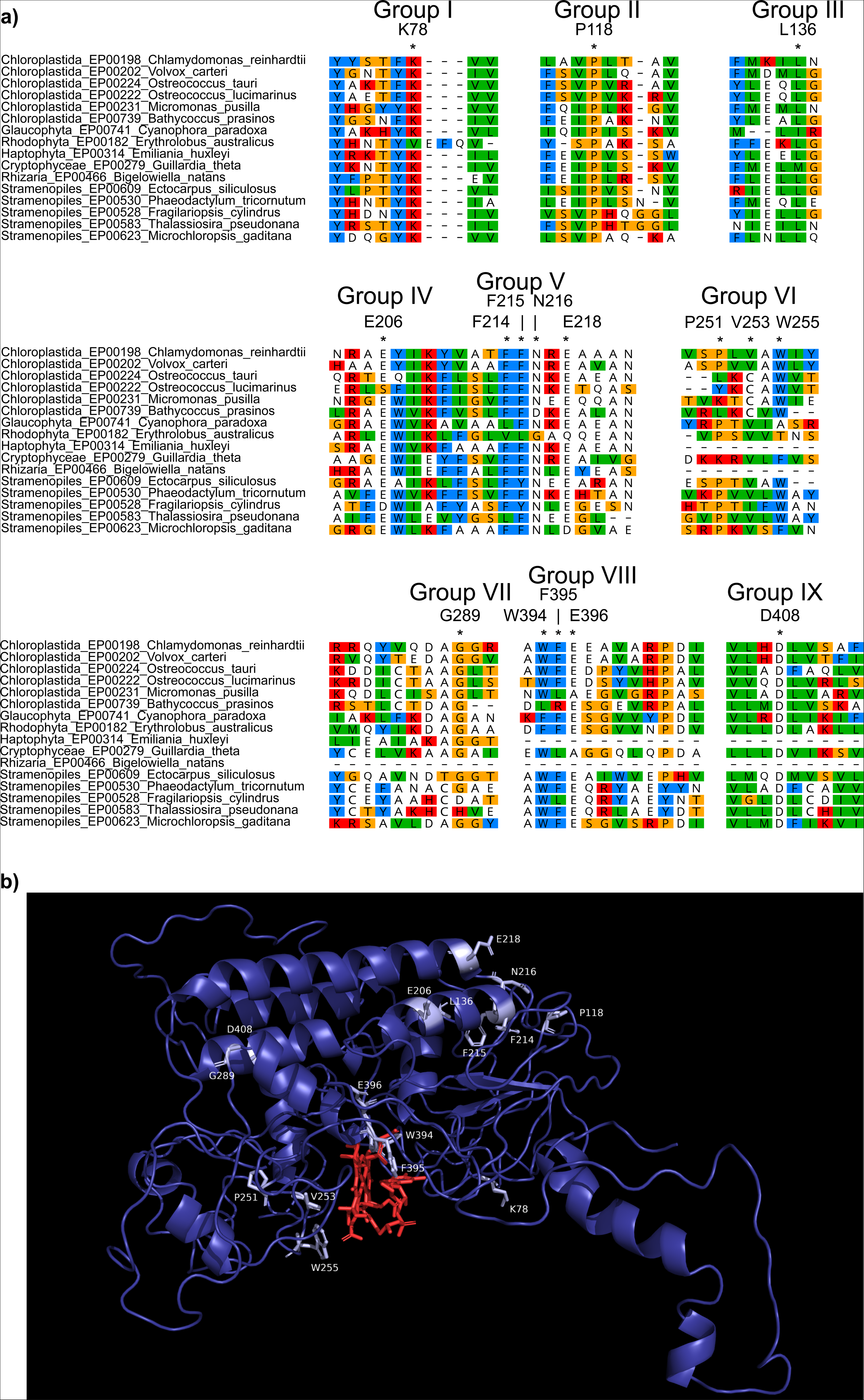
Identification and predicted structural location of CrCBA1 conserved residues. **a)** Sequences with similarity to CBA1 were identified from the EukProt database (Richter et al., 2022) using a manually generated CBA1 Hidden Markov Model (HMM), as described in the materials and methods. A selection of 18 taxa from several eukaryotic supergroups were chosen and conserved regions from the protein are presented. Specific residues indicated by * are: K78, P118, L136, E206, F214, F215, N216, E218, P251, V253, W255, G289, W394, F395, E396 and D408. Protein sequences are coloured according to the Clustal colour-scheme using Geneious Prime 2021.1.1 (www.geneious.com). For each highly conserved region, the corresponding position and amino acid from the CrCBA1 sequence (Cre02.g081050) is indicated. **b)** The predicted 3D structure of CrCBA1 was assessed using the Phyre2 structural prediction server using the intensive mode settings (dark blue). Highly conserved regions of CrCBA1 are indicated in light blue and labelled. CrCBA1 was aligned to the crystal structure of *E. coli* BtuF in complex with B_12_ (pdb: 1n2z). This enabled the relative position of B_12_ (shown in red) to be superimposed onto CrCBA1.

## DISCUSSION

In this study we have shown experimentally that a conserved protein, CBA1, is required for the uptake of the micronutrient B_12_ in two taxonomically distant algae, the diatom *P. tricornutum* (Figure 1) and the chlorophyte *C. reinhardtii* (Figures 3 & 4). Strains with knockouts of the gene were unable to take up B_12,_ demonstrating that there is no functional redundancy of this protein in either organism. This is also the first *in vivo* evidence that CBA1 is present outside the Stramenopiles. Moreover, we found widespread occurrence of CBA1 homologues with considerable sequence conservation across eukaryotic lineages (Figures 7 and S7). The strong correlation of CBA1 with the B_12_-dependent methionine synthase, METH, in algal lineages, provides evidence that CBA1 is a key component of the B_12_ uptake process in evolutionarily distinct microalgae, and the structural similarities between CBA1 and BtuF (Figure 8b), suggest it may operate as a B_12_-binding protein. The highly conserved residues identified in the algal homologues (Figure 8a) offer the means to establish which are functionally important, facilitated by the uptake assay we established.

Nonetheless, the mechanistic role of CBA1 in the process of B_12_ acquisition in algae is not yet clear. Previous physiological studies of B_12_ uptake by microalgae, such as the haptophyte *Diacronema lutheri* (Droop, 1968), indicated a biphasic process: firstly rapid irreversible adsorption of B_12_ to the cell exterior, followed by a slower second step of B_12_ uptake into the cell, consistent with endocytosis. CBA1 is unlikely to be associated with the binding of B_12_ in the cell wall, however. This is because the *C. reinhardtii* strains used in this study, UVM4 and CW15, were cell wall deficient, and therefore likely also deficient in cell wall proteins that bind B_12_; the lack of a B_12_-BODIPY signal from the cell surface in IM4 (Figure S2) supports this hypothesis. Further use of this fluorescent probe offers the possibility to monitor the localisation of B_12_-BODIPIY over time to gain insights into the stages of B_12_ uptake, as has been done in other organisms (Lawrence et al., 2018). In addition, confocal microscopy of CBA1-mVenus fusion protein in *C. reinhardtii* (Figure 5) showed an apparent association of CrCBA1 with the plasma membrane and endomembranes, which is similar to that for ER-localised proteins (Mackinder et al., 2017). Moreover, in a proteomics study of lipid droplets (which form by budding from the ER) CBA1 was in the top 20 most abundant proteins (Goold et al., 2016). Bertrand et al. (2012) found that PtCBA1 had a signal peptide and fluorescently tagged PtCBA1 was also targeted to the ER. Nonetheless, based on its predicted 3D structure and the fact that it has at most one transmembrane helix, CBA1 does not appear to be a transporter itself. Instead, given its structural similarity to BtuF, a distinct possibility is that CBA1 is the soluble component of an ABC transporter, either at the plasma membrane or an internal membrane, and likely will interact with one or more other proteins to allow B_12_ uptake to occur, at least some of them being those involved in receptor-mediated endocytosis, as is the case for B_12_ acquisition in humans (Rutsch et al., 2009; Beedholm-Ebsen et al., 2010; Coelho et al., 2012). In this context, there are known similarities between endocytosis in *C. reinhardtii* and humans (Denning and Fulton, 1989; Bykov et al., 2017), and several putative homologues have been identified by sequence similarity in the alga. Testing the B_12_-uptake capacity of mutants of these proteins would be one approach to investigate whether their roles are also conserved.

In contrast to the situation in algae, the Streptophyta live in a B_12_-free world, neither synthesising nor utilising this cofactor. This is exemplified by the fact that in our analysis only one species, the charophyte alga *Cylindrocystis brebissonii*, encoded METH. Despite this, more than three-quarters of this group encode a CBA1 homologue (Figures 7 & S7). Since the majority of the conserved residues (Figure 8a) are also found in putative CBA1 sequences in the angiosperms such as *Arabidopsis*, including those around the potential binding pocket, it is possible that the streptophyte protein has acquired a new function that still binds a tetrapyrrole molecule. Intriguingly, the reverse is observed in the Metazoa, where METH is almost universal, but CBA1 is entirely absent. However, some Choanoflagellates and some species of fungi do appear to encode both METH and CBA1, suggesting that they utilise B_12_, a trait only recently recognised to occur in fungi (Orłowska et al., 2021). It will be of interest therefore to test whether CBA1 is involved in B_12_ uptake in these organisms, for example by gene knockout studies.

The importance of B_12_ availability for phytoplankton productivity has been demonstrated across several marine ecosystems by amendment experiments (e.g. Bertrand et al., 2011; Koch et al., 2012; Joglar et al., 2021), where addition of B_12_ led to algal blooms and affected the composition and stability of microbial communities. The mode of acquisition of this micronutrient is thus likely to be highly conserved and subject to significant ecological and evolutionary selection pressure to be retained. Moreover, the role of B_12_ at the cellular level may well provide a direct connection between environmental conditions and the epigenetic status of the genome: methionine synthase is the key enzyme in C1 metabolism, linking the folate and methylation cycles and thus responsible for maintaining levels of S-adenosylmethionine (SAM) the universal methyl donor (Hanson & Roje 2001; Mentch & Locasale, 2016). In this context, it is noteworthy that the knockout of *CBA1* in the IM4 line was the result of insertion of a class II transposable element into the gene. This mobilization is likely to reflect epigenetic alterations of the autonomous element, presumably as a result of cellular stress either from the antibiotic selection, or the transformation procedure, or both. Recent classification of the transposons in *C. reinhardtii* indicate that the transposon inserted into *CBA1* in IM4 is a member of the KDZ superfamily of class II TIR elements named Kyakuja-3_cRei (Craig et al. 2021). If the phenomenon of inactivation of a gene that is deleterious (in this case allowing B_12_ to be taken up and repress the antibiotic resistance gene) via transposition is a general response in *C. reinhardtii*, repeating the screen for CBA1 mutants might allow observation of further transposition events, and enable characterisation of this group of elements at the functional level. Moreover, it could be adopted as a more general methodology to identify candidate genes involved in other physiological processes, by tying their expected effects to deleterious outcomes through synthetic biology constructs and screening surviving mutants by sequencing.

## MATERIALS AND METHODS

### Organisms and growth conditions

Strains, media and growth conditions used in this study are listed in Table S1. If required, antibiotics, vitamin B_12_ (cyanocobalamin) and thiamine were added to the medium at concentrations indicated. Algal culture density was measured using a Z2 particle count analyser (Beckman Coulter Ltd.) and optical density (OD) at 730 nm was measured using a FluoStar OPTIMA (BMG labtech) plate reader or a CLARIOstar plate reader (BMG labtech). Bacterial growth was recorded by measuring OD_595_.

### Algal B_12_-uptake assay

Algal cultures were grown to stationary phase and cyanocobalamin salt (Sigma) was added (*P. tricornutum*: 600 pg; *C. reinhardtii*: 150 pg) to 5 x 10^6^ cells in a final volume of 1 ml in f/2 or TAP medium respectively. The samples were incubated at 25°C under continuous light with shaking for 1 hour and inverted every 30 minutes to aid mixing. Samples were centrifuged and the supernatant (media fraction) transferred into a fresh microcentrifuge tube. The cell pellet was resuspended in 1 ml water. Both samples were boiled for 10-20 minutes to release any cellular or bound B_12_ into solution, and then centrifuged to pellet debris. The supernatant was used in the *S. typhimurium* B_12_ bioassay as described in Bunbury et al. (2020). The amount of B_12_ in the sample was calculated by comparison to a standard curve of known B_12_ concentrations fitted to a 4 parameter logistic equation f(x) = c + (d-c)(1+exp(b(log(x)-log(e)))) (Ritz et al., 2015). This standard curve was regenerated with every bioassay experiment.

### Generating *P. tricornutum* CBA1 knockout lines using CRISPR-Cas9

CRISPR/Cas9 genome editing applied the single guide RNA (sgRNA) design strategy described in Hopes et al., (2017). Details are provided in the Supplementary methods. *P. tricornutum CCAP 1055/1* cells were co-transformed with linearised plasmids pMLP2117 and pMLP2127 using a NEPA21 Type II electroporator (Nepa Gene) as previously described (Yu et al., 2021). After plating on 1% agar selection plates containing 75 mg·l^−1^ zeocin and incubation for 2-3 weeks, zeocin resistant colonies were picked into 96 well plates containing 200 µl of f/2 media with 75 mg·l^−1^ zeocin. After seven days strains were subcultured into fresh media either containing 75 mg·l^−1^ zeocin or 300 mg·l^−1^ nourseothricin, and genotyped with a three-primer PCR using PHIRE polymerase (Thermo Fisher Scientific) with primers gCBA1.fwd, gCBA1.rv and NAT.rv (Table S2). Five promising colonies resistant to nourseothricin and with genotypes showing homologous recombination or indels were re-streaked on 75 mg·l^−1^ zeocin f/2 plates to obtain secondary monoclonal colonies. Twelve secondary colonies were picked for each primary colony after 2-3 weeks and again genotyped with a three-primer PCR. Promising colonies were genotyped in further detail with primer pairs gCBA1.fwd/gCBA1.rv, gCBA1.fwd/NAT.rv and gCBA1in.fwd/gCBA1in.rv (Table S2).

### Construct assembly and *C. reinhardtii* transformation

Constructs were generated using Golden Gate cloning, using parts from the *Chlamydomonas* MoClo toolkit (Crozet et al., 2018) and some that were created in this work. All parts relating to *Cre02.g081050* were domesticated from UVM4 genomic DNA, with BpiI and BsaI sites removed from the promoter, ORF and terminator by PCR based mutagenesis. A list of plasmids used in this study is shown in Table S2. Transformation of *C. reinhardtii* cultures with linearised DNA was carried out by electroporation essentially as described by Mehrshahi et al. (2020) before plating on TAP-agar plates with the appropriate antibiotics.

Insertional mutagenesis was performed as above, however, cultures were grown to a density of approximately 1×10^7^ cells/ml and were incubated with 500 ng transgene cassette. After allowing the cells to recover overnight in TAP plus 60 mM sucrose at 25°C in low light (less than 10 μmol photon m^−2^.s^−1^ at 100 rpm), between 200 - 250 μl of transformants were plated on solid TAP media (square 12×12 cm petri dishes) containing ranges of 15-20 μg/ml hygromycin, 20-50 μg/ml paromomycin and 48-1024 ng/l vitamin B12, and the plates were incubated in standing incubators.

### Confocal laser scanning microscopy

*C. reinhardtii* transformants carrying the pAS_C2 construct were imaged in a confocal laser scanning microscope (TCS SP8, Leica Microsystems, Germany) with an HC PL APO CS2 40x/1.30 aperture oil-immersion lens. Images were taken using the sequential mode provided by the Leica LAS software, with the channel used for mVenus and brightfield detection being taken first with excitation from a white light source at 486 nm and emissions were detected between 520 - 567 nm, followed by chlorophyll detection (excitation 514 nm, emission 687-724 nm). The overlay images were produced automatically by the Leica LAS software. Inkscape was used to increase the lightness and decrease the contrast of all the images in the same manner.

### Quantitative real-time PCR

Quantification of steady state levels of transcripts was carried out according to Bunbury et al. (2020), using random hexamer primers for cDNA synthesis. The qPCR data was analysed using the ΔΔCT method with an assumed amplification efficiency of 2. Log2(2-ΔCT) values were plotted in the resulting figures.

### Whole genome sequencing

Genomic DNA was extracted from *C. reinhardtii* cells by phenol-chloroform extraction and sequenced using the NovaSeq sequencing platform by Novogene (Cambridge, UK) to produce 150 bp paired-end reads. This involved RNase treatment and library preparation with the NEBNext Ultra II DNA Library Prep Kit (PCR-free), which generated 350 bp inserts. The raw sequencing data for this study have been deposited in the European Nucleotide Archive (ENA) at EMBL-EBI under accession number PRJEB58730 (https://www.ebi.ac.uk/ena/browser/view/PRJEB58730). Novogene performed all quality filtering, summary statistics and bioinformatic analysis. The location of the Hyg3 cassette was determined by identifying loci that comprised reads from IM4 that mapped between genomic DNA and pHyg3, and cross-referencing these loci against the parental strains. The TE identification was carried out similarly, full details are provided in Supplementary Methods.

### Bioinformatics pipeline

The EukProt database was assessed for the presence of METE, METH and CBA1 (Richter et al., 2022). The query used for CBA1 was a hidden Markov model (HMM) generated from the protein fasta sequences: Phatr3_J48322, Thaps3 11697, Fracy1 241429, Fracy1 246327, Auran1 63075, Ectocarpus siliculosus D8LMT1 and Cre02.g081050.t1.2 by first aligning using MAFFT (Katoh and Standley, 2013) version 7.470 with the --auto option, and then building a HMM using hmmbuild (hmmer 3.2.1). Additionally, protein fasta (Cre06.g250902, Cre03.g180750), PFAM (PF02310, PF02965, PF00809, PF02574, PF01717, PF08267) and KO (K00548, K00549) queries were searched against EukProt to identify sequences with similarity to METE and METH. The queries were searched against EukProt using hmmsearch (HMMER 3.1b2). The default bitscore thresholds were used for KO and PFAM queries. The threshold used for CBA1 HMM, and the CrMETE and CrMETH protein fasta sequences, was a full-length e-value of 1e-20. For each protein, all individual queries were required to be significant to classify the protein as present. The best hit in each species was identified by taking the protein with the greatest geometric mean of full length bitscores for the queries. The dataset was joined with taxonomic information from EukProt and completeness information calculated using BUSCO version 4.1.4 and eukaryote_odb10 (Manni et al., 2021).

## Supporting information

Sayer et al. Supplementary material

## Acknowledgements

We thank Catherine Sutherland for help with maintaining and screening the CRISPR/Cas9 mutants of *P. tricornutum* and Dr Lorraine Archer for lab management. We are grateful to Dr Amanda Hopes and Prof Thomas Mock (University of East Anglia) for the parts used in gene editing of *PtCBA1*. The plasmid Hyg3 used in the insertional mutagenesis was obtained from the Chlamydomonas Resource Centre (www.chlamycollection.org). This work was supported by the UK’s Biotechnology and Biological Sciences Research Council (BBSRC) Doctoral Training Partnership, grant no. BB/M011194/1 to A.P.S., M.L.P. and A.G.S; grant no. BB/M018180/ 1 to P.M. and A.G.S.; grant no. BB/L002957/1 and BB/R021694/1 to K.G. and AGS; grant no. BB/L014130/1 to G.I.M, K.G., P.M. and A.G.S; grant no. BB/S002197/1 to M.J.W; University of Cambridge Broodbank Fellowship to G.I.M; Royal Society grant no. INF\R2\180062 to M.J.W.; and Bill & Melinda Gates Foundation grant OPP1144 and Gates Cambridge Trust (Graduate Student Fellowship) to A.H. For the purpose of open access, the authors have applied a Creative Commons Attribution (CC BY) licence to any Author Accepted Manuscript version arising from this submission.

## Author contributions

APS designed and performed research, analysed data and wrote the article with contributions from all the authors. KG and MLP carried out the CRISPR/Cas9 editing of Phaeodactylum and contributed to writing the article. AH carried out the bioinformatics analysis to identify the putative transposable elements. MJW & ADL synthesised the BODIPY-labelled B_12_ and contributed to writing the article. KG, GMO and PM supervised aspects of the project and contributed to writing the article. AGS conceived the project, obtained the funding, supervised the project and wrote the article with contributions from all the authors. AGS agrees to serve as the author responsible for contact and ensures communication.

---

**Figure S1.**
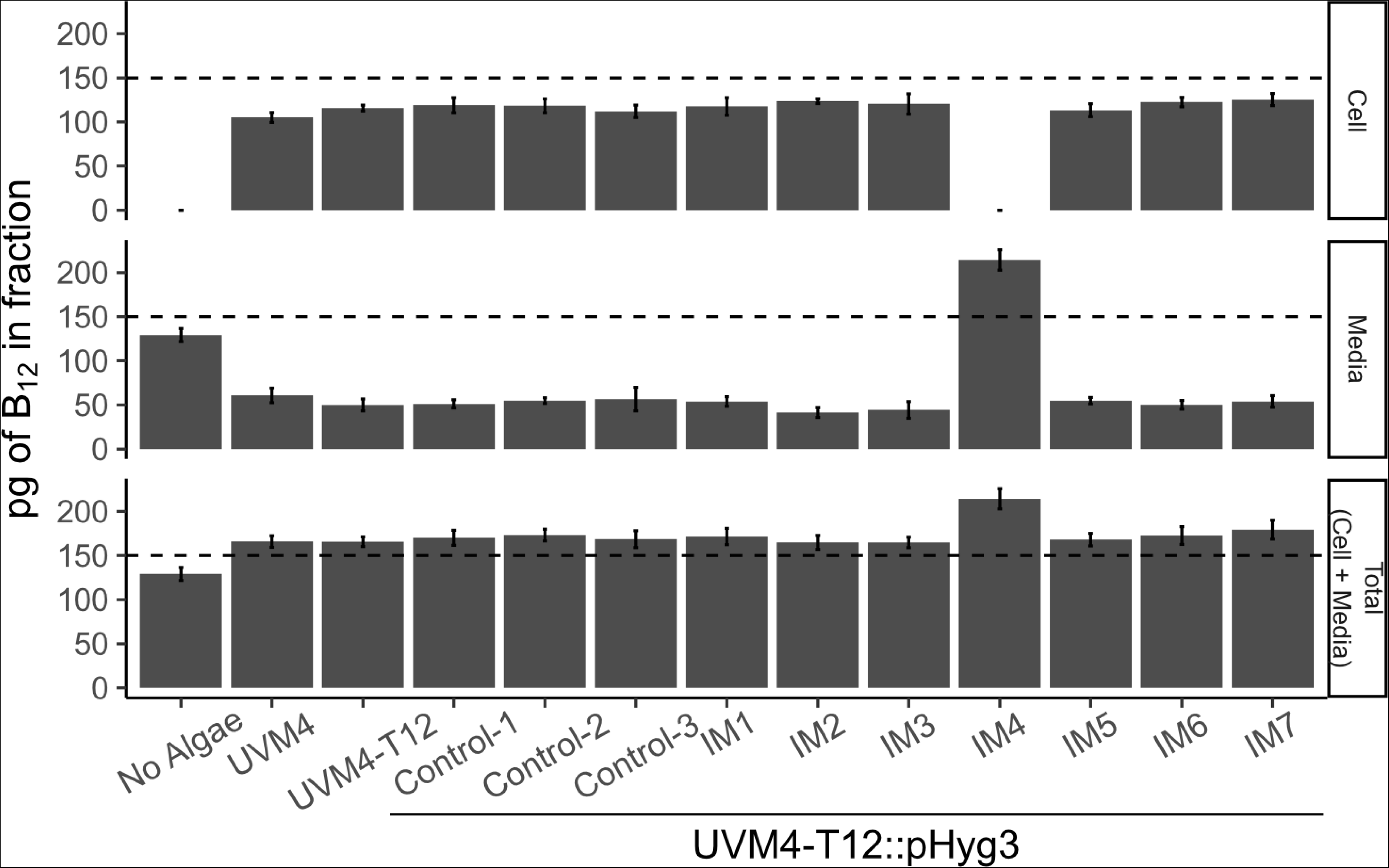
Characterisation of B12 uptake in *C. reinhardtii* insertional mutant lines. To determine whether any of the 7 insertional mutant lines (IM) isolated from mutagenesis of UVM4::T12 showed impaired B12 uptake, the B12-uptake assay was performed as described in the materials and methods. Control-1, Control-2 and Control-3 lines were picked from solid media with paromomycin and hygromycin but without vitamin B12, whereas IM1-IM7 lines were picked from solid media containing paromomycin, hygromycin and vitamin B12. The total was inferred by the addition of the cell and media fractions. The dashed line indicates the amount of B12 added in the uptake assay. Standard deviation error bars are shown, n=4. Statistical analysis was performed on the media fraction, and Tukey’s test identified the following comparisons to be significantly different from one another (only reporting IM strains different from UVM4, UVM4-T12 or Control-[1,2,3] strains): IM4 vs UVM4 (p<1e^−12^), UVM4-T12 (p<1e^−12^), Control-1 (p<1e^−12^), Control-2 (p<1e^−12^), Control-3 (p<1e^−12^); and IM2 vs UVM4 (p<0.05).

**Figure S2.**
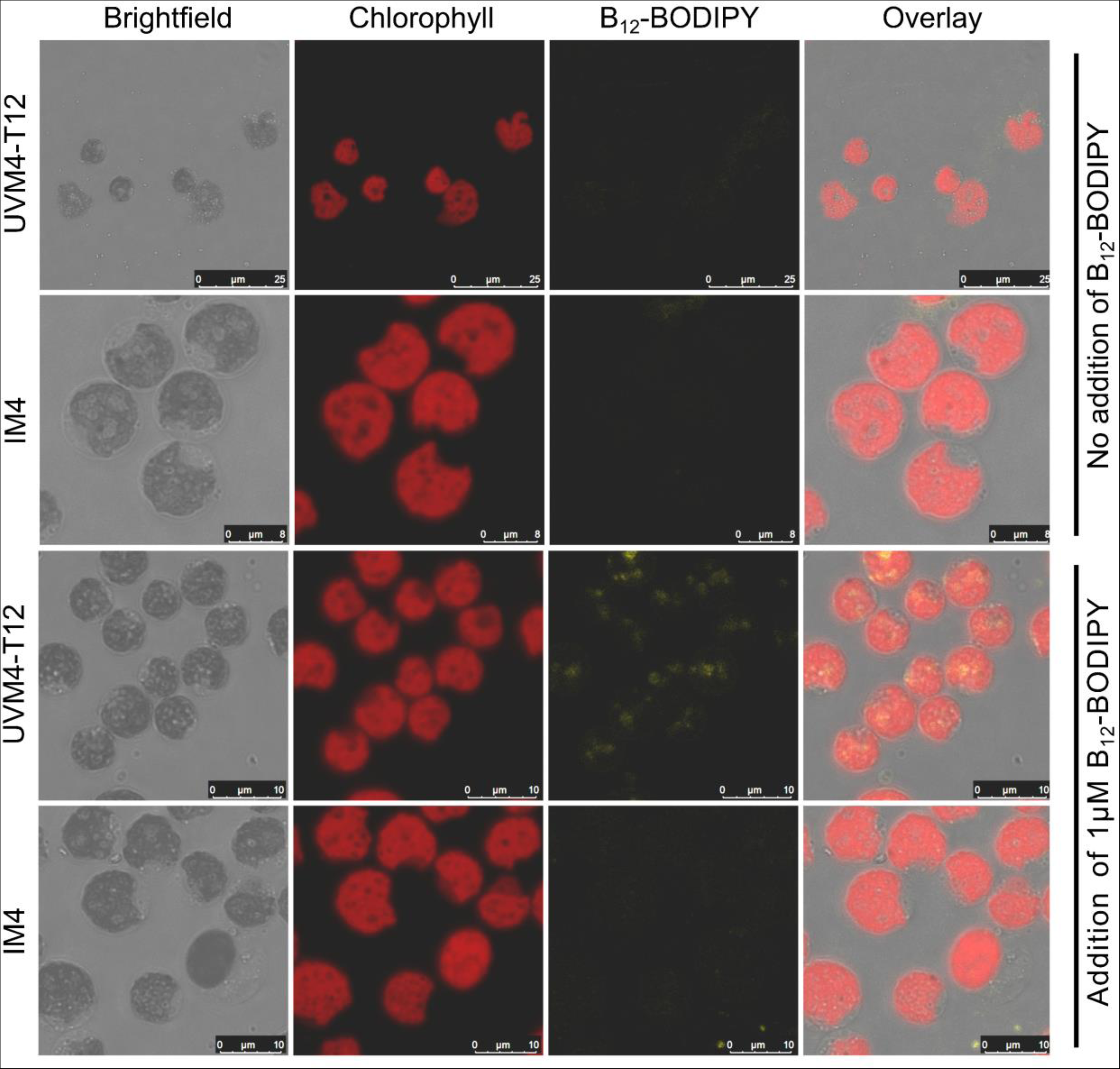
Visualisation of B12-BODIPY uptake in *C. reinhardtii* using confocal microscopy. To assess B12 uptake more directly, the insertional mutant line IM4 (impaired B12 uptake) and its parental line UVM4-T12 were incubated with the fluorescent B12 analogue B12-BODIPY and the samples were imaged using confocal microscopy, as described in the materials and methods. Channels shown are brightfield (greyscale), chlorophyll (red), B12-BODIPY (yellow) and an overlay.

**Figure S3.**
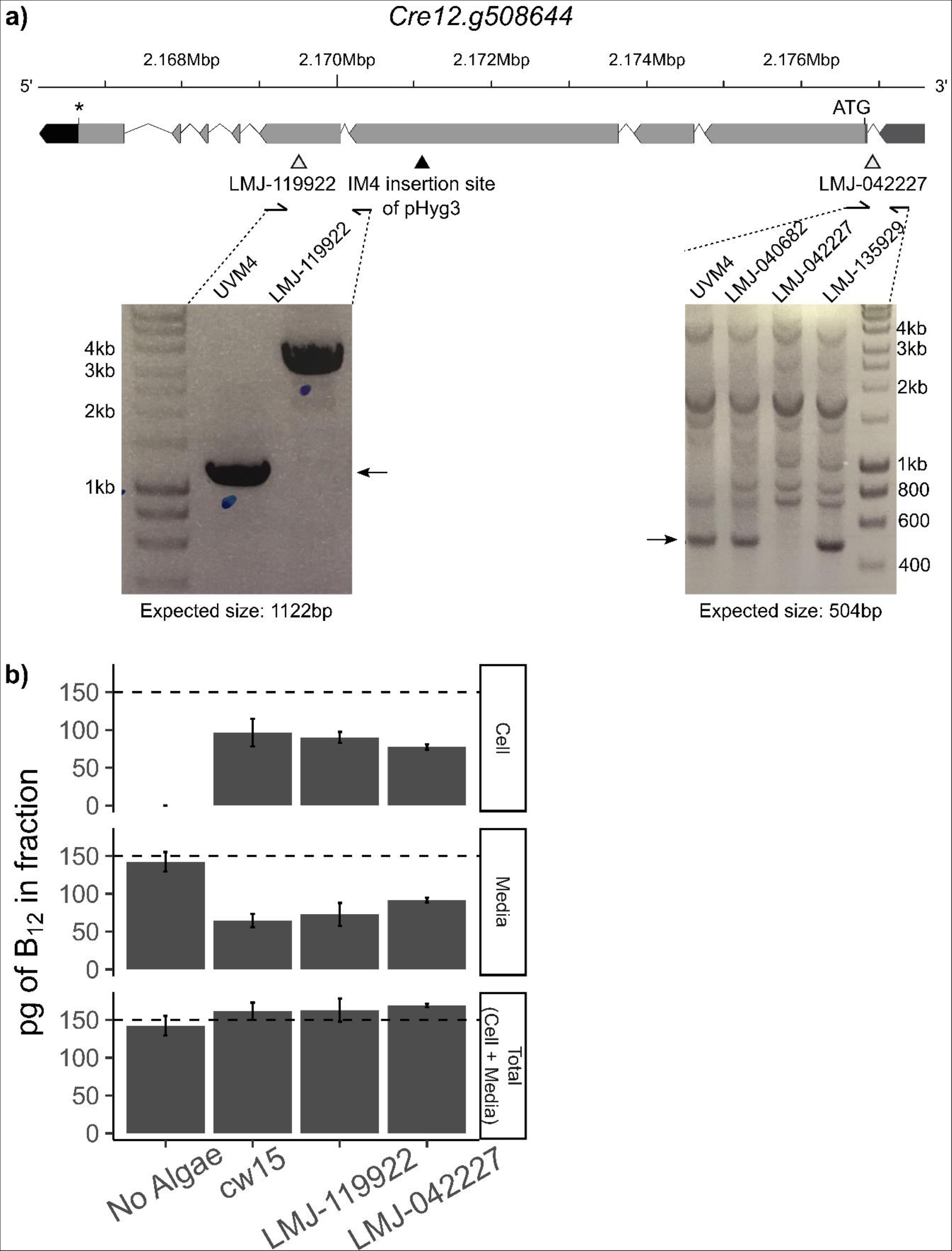
*C. reinhardtii* knockout lines of *Cre12.g508644* are able to take up B12. **a)** Schematic showing the structure of *Cre12.g508644* with annotations for the 5’UTR (medium grey), start codon (ATG), exons (light grey), introns (black lines), stop codon (*) and 3’UTR (black). The location of the IM4 pHyg3 insertion site (identified by DNA sequencing and validated using PCR) is indicated with a black triangle. The predicted disruption sites in CLiP (Li et al., 2016) knockout strains LMJ-119922 and LMJ- 042227 are indicated with grey triangles. These were also confirmed by PCR (insets) **b)** B12-uptake assay of the CLiP mutants and their background strain cw15. The dashed line shows the amount of B12 added to the experiment. Total = inferred by addition of amounts determined in the cellular and media fractions. Standard deviation error bars are shown, No Algae (n=10), cw15 (n=10), LMJ-119922 (n=10) and LMJ- 042227 (n=6). Statistical analysis was performed on the media fraction, and Tukey’s test identified the following comparisons to be significantly different from one another: No Algae vs cw15 (p<1e-12); No Algae vs LMJ-119922 (p<1e-12); No Algae vs LMJ-042227 (p<1e-08); cw15 vs LMJ-042227 (p<1e-03); and LMJ-119922 vs LMJ-042227 (p<0.05).

**Figure S4.**
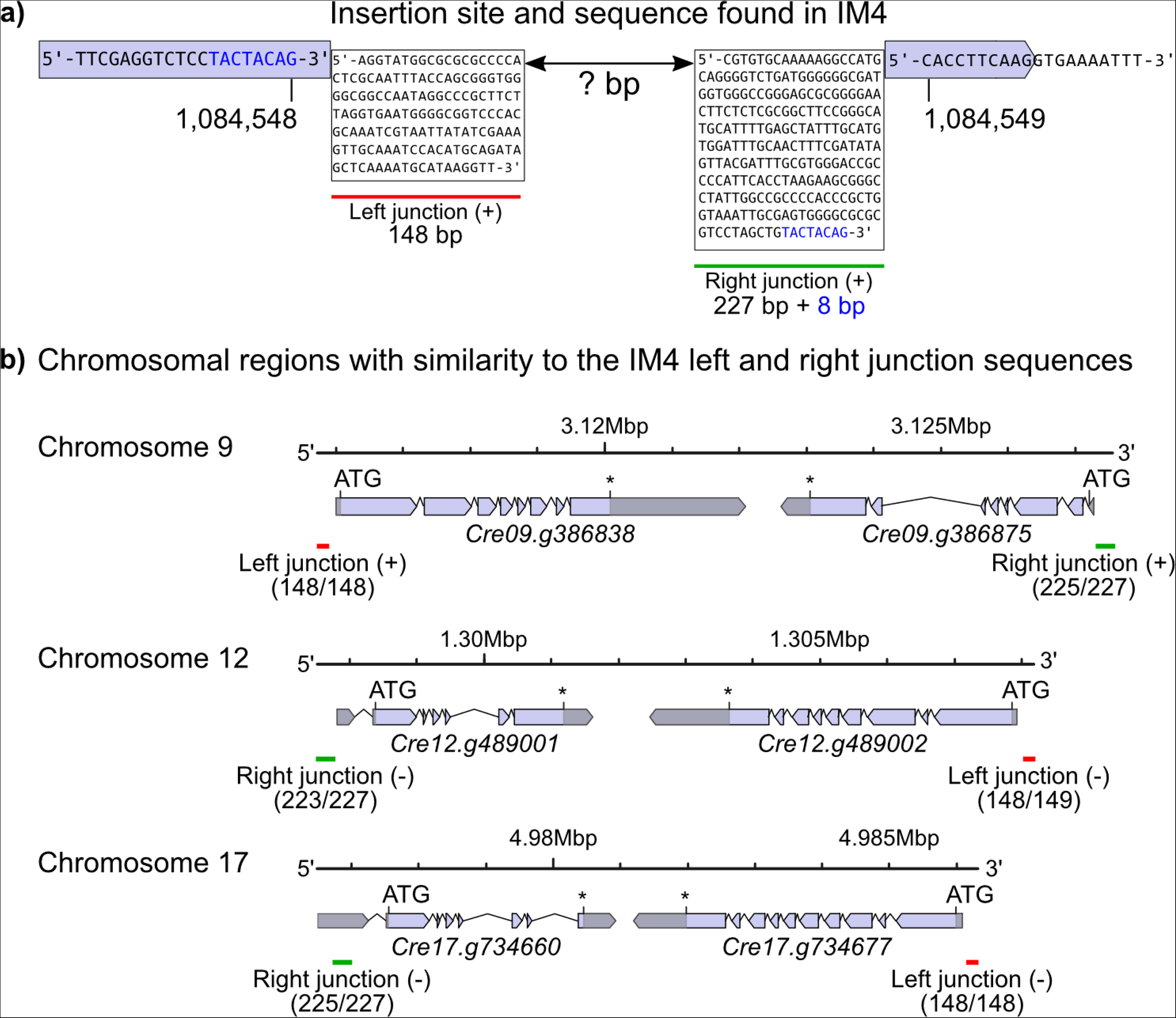
Structure and sequence of a second insertion in the IM4 strain. **a)** Mapping of the WGS data to the reference strain revealed an extra sequence between positions 1,084,548 bp and 1,084,549 bp of chromosome 2 in the *C. reinhardtii* v5.0 Phytozome reference genome corresponding to the Cre02.g081050 locus. An 8 bp target site duplication ‘TACTACAG’ was identified flanking the insertion (blue). The sequences of the left (red) and right (green) insertion junctions were identified from DNA sequencing reads and confirmed with PCR and sequencing. The sequence between these left and right junctions has not been determined. **b)** Chromosomal regions with similarity to the left (148 bp) and right (227 bp) junction sequences were identified by a BLAST search. Shown are three regions with sequence similarity to the left and right junctions and where the left and right junctions are a similar distance apart (~10 Kb) and in the same orientation. The position of the matching left junction (red) and right junction (green) sequences are indicated. The matching strand is indicated in brackets. The number of matching base pairs from the BLAST search is listed in brackets (target/query).

**Figure S5.**
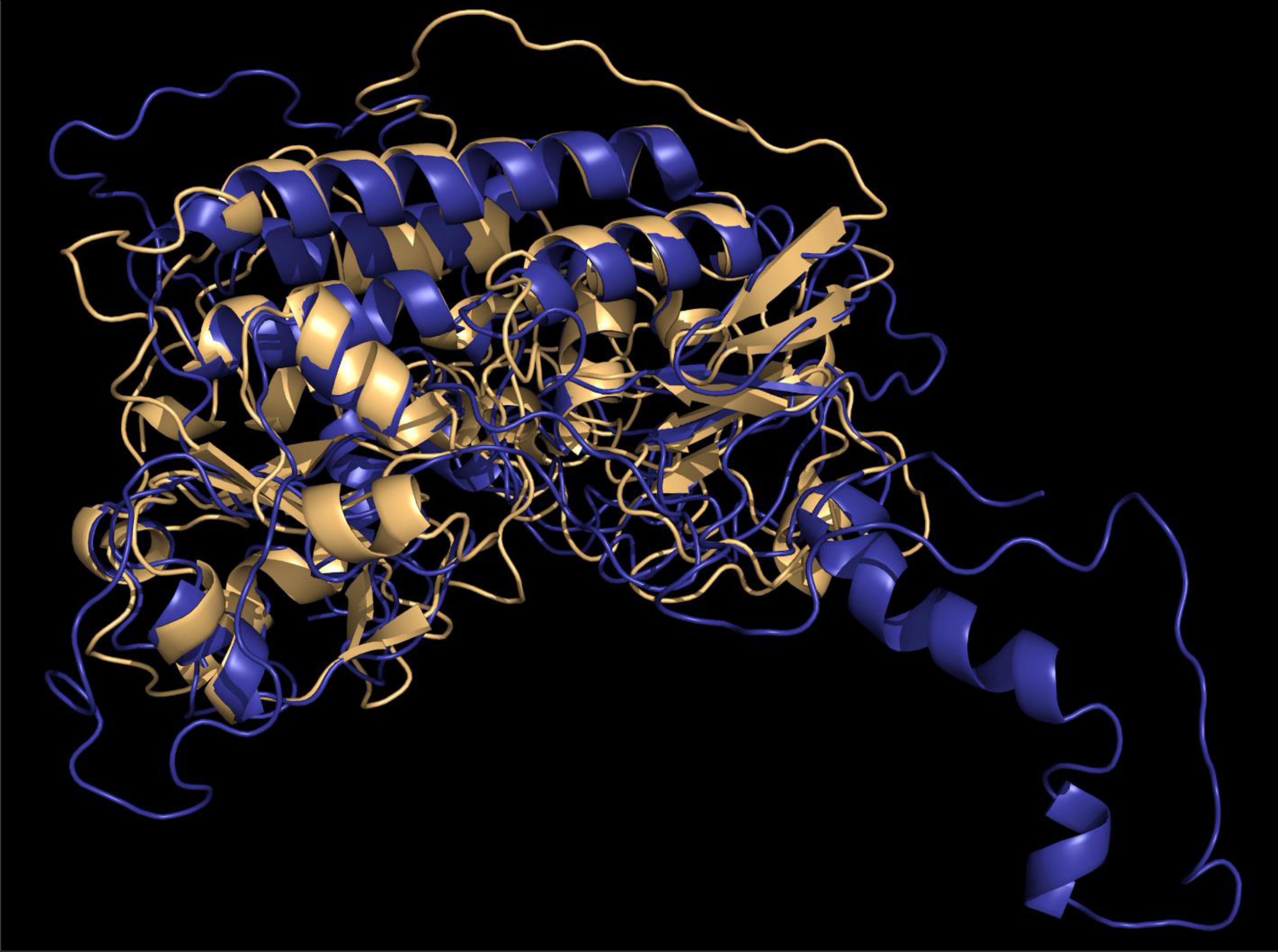
The predicted structures of CrCBA1 and PtCBA1 show a high degree of structural similarity. Structural prediction of CrCBA1 (Cre02.g081050) (blue) and PtCBA1 (Phatr3_J48322) (gold) was performed using the Phyre2 structural prediction server (Kelley et al., 2015). CrCBA1 was modelled with 56% of residues predicted with greater than 90% accuracy; PtCBA1 was modelled with 69% of residues predicted with greater than 90% accuracy. Structures were aligned using the super command in PyMOL, root mean squared deviation = 2.333. Conserved alpha helices are seen in the centre of the image, as is a common cleft region at the bottom of the image.

**Figure S6.**
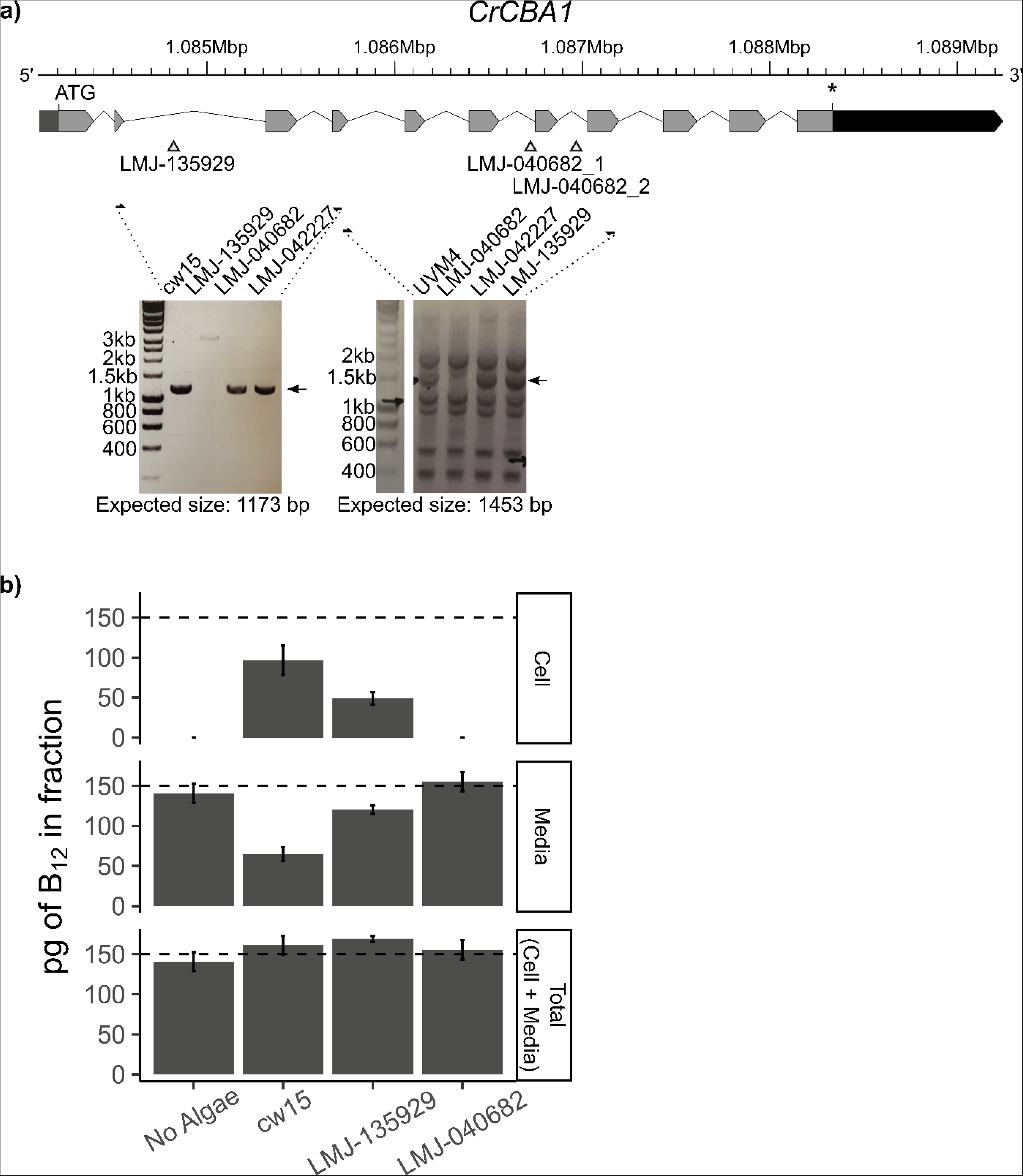
Independent mutant lines of *CrCBA1* show defective B12 uptake. **a)** Schematic showing the structure of *CrCBA1* (Cre02.g081050) with annotations for the 5’UTR (medium grey), start codon (ATG), exons (light grey), introns (black lines), stop codon (*) and 3’UTR (black). The location of predicted disruption sites in *CrCBA1* CliP knockout strains LMJ-135929 and LMJ-040682 are indicated (triangles). Primer positions used for the analysis of mutant lines are shown with arrows. Insets show that primers amplifying between exons 2 and 4 produced a band of the expected size for control lines cw15, LMJ- 040682 and LMJ-042227, whereas LMJ-135929 showed an increased amplicon size, indicating an insertion in this region. Primers between exons 4 and 8 produced a band of the expected size in UVM4 and the same-background control lines LMJ-042227 and LMJ-135929, whereas LMJ-040682 lacked this band, indicating a disruption in this region. **B)** B12-uptake assay. The dashed line indicates the amount of B12 added to the sample. Standard deviation error bars are shown, No Algae (n=14), cw15 (n=10), LMJ- 135929 (n=6) and LMJ-040682 (n=18). Statistical analysis was performed on the media fraction, and Tukey’s test identified the following comparisons to be significantly different from one another: No Algae vs cw15 (p<1e-12); No Algae vs LMJ-135929 (p<1e-02); No Algae vs LMJ-040682 (p<1e-02); cw15 vs LMJ- 135929 (p<1e-11); cw15 vs LMJ-040682 (p<1e-12); and LMJ-135929 vs LMJ-040682 (p<1e-06).

**Figure S7.**
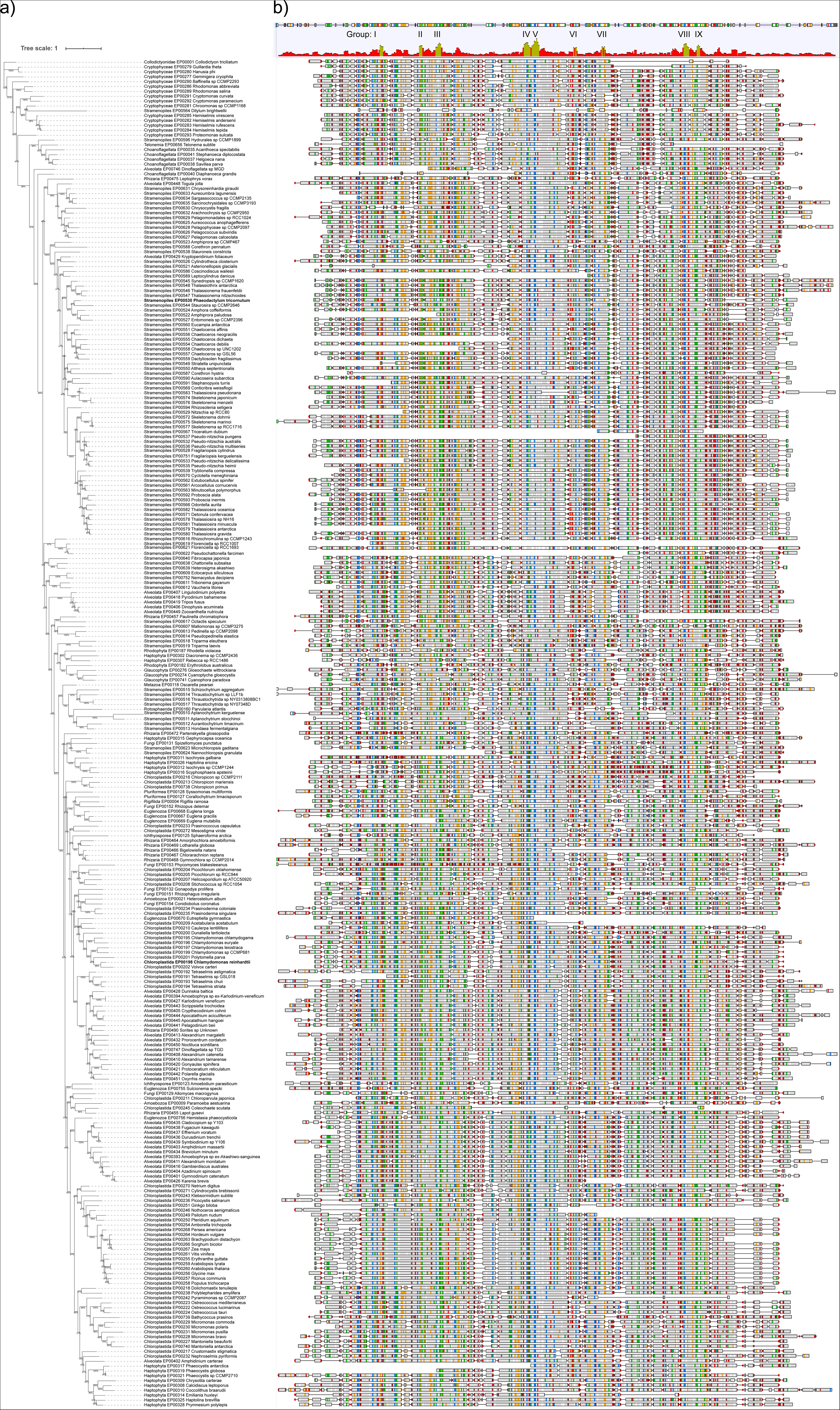
Sequences with similarity to CBA1 are found throughout Eukaryota. Sequences with similarity to CBA1 were identified from the EukProt database (Richter et al., 2022) using a manually generated hidden Markov model for CBA1, as described in the materials and methods. Positively classified CBA1 sequences were aligned with MAFFT (-- auto option) (Katoh and Standley, 2013) and trimmed with trimai (-automated1 option) (Capella-Gutiérrez et al., 2009), and Iqtree version 1.6.10 (options: -bb 1000, -safe, -bnni, - alrt 1000, -st AA, -seed 1000, -msub nuclear, -t RANDOM, -m TEST) (Nguyen et al., 2015) was used to produce a gene tree. Visualisation was performed using iTOL (Letunic and Bork, 2019). A multiple sequence alignment of the full length protein sequences was generated and visualised using Geneious Prime 2021.1.1 (https://www.geneious.com), and is shown adjacent to the tree (options: Clustal colour scheme, agreements to consensus highlighted, sliding window size = 5 bp, and sites with over 50% gaps hidden). A conservation track and consensus track are shown above the alignment panel, with regions of conserved residues indicated, detailed more in Figure 8a.

## References

Allen, MD, del Campo JA, Kropat J, Merchant SS (2007) FEA1, FEA2, and FRE1, encoding two homologous secreted proteins and a candidate ferrireductase, are expressed coordinately with FOX1 and FTR1 in iron-deficient *Chlamydomonas reinhardtii*. Eukaryotic Cell 6: 1841–1852

Almagro Armenteros J, Sønderby CK, Sønderby SK, Nielsen H, Winther O (2017) DeepLoc: Prediction of protein subcellular localization using deep learning. Bioinformatics 33: 3387–3395

Almagro Armenteros J, Tsirigos KD, Sønderby CK, Petersen TN, Winther O, Brunak S, Heijne G von, Nielsen H (2019) SignalP 5.0 improves signal peptide predictions using deep neural networks. Nature Biotechnology 37: 420–423

Banerjee R, Gouda H, Pillay S (2021) Redox-linked coordination chemistry directs vitamin B_12_ trafficking. Accounts of Chemical Research 54: 2003–2013

Beedholm-Ebsen R, Wetering KVD, Hardlei T, Nexø E, Borst P, Søren K, Moestrup SK (2010) Identification of multidrug resistance protein 1 (MRP1/ABCC1) as a molecular gate for cellular export of cobalamin. Blood 115: 1632–1639

Bertrand EM, Saito MA, Rose JM, Riesselman CR, Lohan MC, Noble AE, Lee PA, DiTullio GR (2011) Vitamin B_12_ and iron colimitation of phytoplankton growth in the Ross Sea. Limonol Oceanogr 52: 1079–1093

Bertrand EM, Allen AE, Dupont CL, Norden-Krichmar TM, Bai J, Valas RE, Saito MA (2012) Influence of cobalamin scarcity on diatom molecular physiology and identification of a cobalamin acquisition protein. Proc Natl Acad Sci U S A 109: E1762–71

Borths EL, Locher KP, Lee AT, Rees DC (2002) The structure of *Escherichia coli* BtuF and binding to its cognate ATP binding cassette transporter. Proc Natl Acad Sci U S A 99: 16642–7

Bunbury F, Helliwell KE, Mehrshahi P, Davey MP, Salmon DL, Holzer A, Smirnoff N, Smith AG (2020) Responses of a newly evolved auxotroph of Chlamydomonas to B_12_ deprivation. Plant Physiology 183: 167–178

Bykov YS, Schaffer M, Dodonova SO, Albert S, Plitzko JM, Baumeister W, Engel BD, Briggs JA (2017) The structure of the COPI coat determined within the cell. eLife 6: e32493

Carlucci FA, Silbernagel BS, McNally MP (2007) The influence of temperature and solar radiation on persistence of vitamin B12, thiamine, and biotin in seawater. Journal of Phycology 5: 302–305

Choi CC, Ford RC (2021) ATP binding cassette importers in eukaryotic organisms. Biological Reviews 96: 1318–1330

Coelho D, Kim JC, Miousse IR, Fung S, Moulin M du, Buers I, Suormala T, Burda P, Frapolli M, Stucki M, et al (2012) Mutations in ABCD4 cause a new inborn error of vitamin B12 metabolism. Nature Genetics 44: 1152–1155

Craig RJ, Hasan AR, Ness RW, Keightley PD (2021) Comparative genomics of *Chlamydomonas*. Plant Cell 33:1016–1041

Croft MT, Lawrence AD, Raux-Deery E, Warren MJ, Smith AG (2005) Algae acquire vitamin B_12_ through a symbiotic relationship with bacteria. Nature 438: 90–3

Crozet P, Navarro FJ, Willmund F, Mehrshahi P, Bakowski K, Lauersen KJ, Pérez-Pérez M-E, Auroy P, Gorchs Rovira A, Sauret-Gueto S, et al (2018) Birth of a photosynthetic chassis: A MoClo toolkit enabling synthetic biology in the microalga *Chlamydomonas reinhardtii*. ACS Synthetic Biology 7: 2074–2086

Denning GM, Fulton AB (1989) Purification and characterization of clathrin-coated vesicles from *Chlamydomonas*. The Journal of Protozoology 36: 334–340

Droop MR (1968) Vitamin B12 and marine ecology. IV. The kinetics of uptake, growth and inhibition in Monochrysis lutheri. Journal of the Marine Biological Association of the United Kingdom 48: 689–733

Field CB, Behrenfeld MJ, Randerson JT, Falkowski P (1998) Primary production of the biosphere: Integrating terrestrial and oceanic components. Science 281: 237–240

Gonzalez JC, Banerjee RV, Huang S, Sumner JS, Matthews RG (1992) Comparison of cobalamin-independent and cobalamin-dependent methionine synthases from *Escherichia coli*: Two solutions to the same chemical problem. Biochemistry 31: 6045–6056

Goold HD, Cuiné S, Légeret B, Liang Y, Brugière S, Auroy P, Javot H, Tardif M, Jones B, Beisson F, et al (2016) Saturating light induces sustained accumulation of oil in plastidal lipid droplets in *Chlamydomonas reinhardtii*. Plant Physiology 171: 2406–2417

Hanson AD, Roje S (2001) One-Carbon metabolism in higher plants. Ann Rev Plant Physiol 52: 119–137

Helliwell KE, Collins S, Kazamia E, Purton S, Wheeler GL, Smith AG (2015) Fundamental shift in vitamin B_12_ eco-physiology of a model alga demonstrated by experimental evolution. ISME J 9: 1446–1455

Helliwell KE, Scaife MA, Sasso S, Araujo APU, Purton S, Smith AG (2014) Unraveling vitamin B_12_-responsive gene regulation in algae. Plant Physiol 165: 388–397

Helliwell KE, Wheeler GL, Leptos KC, Goldstein RE, Smith AG (2011) Insights into the evolution of vitamin B_12_ auxotrophy from sequenced algal genomes. Molecular Biology and Evolution 28: 2921–2933

Hopes A, Nekrasov V, Belshaw N, Grouneva I, Kamoun S, Mock T (2017) Genome editing in diatoms using CRISPR-cas to induce precise bi-allelic deletions. Bio Protoc 7: e2625

Joglar V, Pontiller B, Martínez-García S, Fuentes-Lema A, Pérez-Lorenzo M, Lundin D, Pinhassi J, Fernández E, Teira E (2021) Microbial plankton community structure and function responses to vitamin B_12_ and B_1_ amendments in an upwelling system. Appl Environ Microbiol 87: e0152521.

Kadner RJ (1990) Vitamin B_12_ transport in *Escherichia coli*: Energy coupling between membranes. Molecular Microbiology 4: 2027–2033

Katoh K, Standley DM (2013) MAFFT multiple sequence alignment software version 7: Improvements in performance and usability. Molecular Biology and Evolution 30: 772–780

Kelley LA, Mezulis S, Yates CM, Wass MN, Sternberg MJE (2015) The Phyre2 web portal for protein modeling, prediction and analysis. Nat Protoc 10: 845–58

King N, Westbrook MJ, Young SL, Kuo A, Abedin M, Chapman J, et al. (2008). The genome of the choanoflagellate *Monosiga brevicollis* and the origin of metazoans. Nature. 451: 783–788.

Koch F, Hattenrath-Lehmann TK, Goleski JA, Sañudo-Wilhelmy S, Fisher NS, Gobler CJ. 2012. Vitamin B_1_ and B_12_ uptake and cycling by plankton communities in coastal ecosystems. Front Microbiol 3:363.

Lawrence AD, Nemoto-Smith E, Deery E, Baker JA, Schroeder S, Brown DG, Tullet JMA, Howard MJ, Brown IR, Smith AG, et al (2018) Construction of fluorescent analogs to follow the uptake and distribution of cobalamin (vitamin B_12_) in bacteria, worms, and plants. Cell Chem Biol 25: 941–951

Li X, Zhang R, Patena W, Gang SS, Blum SR, Ivanova N, Yue R, Robertson JM, Lefebvre PA, Fitz-Gibbon ST, et al (2016) An indexed, mapped mutant library enables reverse genetics studies of biological processes in *Chlamydomonas reinhardtii*. Plant Cell 28: 367–87

Mackinder LCM, Chen C, Leib RD, Patena W, Blum SR, Rodman M, Ramundo S, Adams CM, Jonikas MC (2017) A spatial interactome reveals the protein organization of the algal CO_2_ concentrating mechanism. Cell 171: 133–147.e14

Manni M, Berkeley MR, Seppey M, Simão FA, Zdobnov EM (2021) BUSCO update: Novel and streamlined workflows along with broader and deeper phylogenetic coverage for scoring of eukaryotic, prokaryotic, and viral genomes. Molecular Biology and Evolution 38: 4647–4654

Mehrshahi P, Nguyen GTDT, Gorchs Rovira A, Sayer A, Llavero-Pasquina M, Lim Huei Sin M, Medcalf EJ, Mendoza-Ochoa GI, Scaife MA, Smith AG (2020) Development of novel riboswitches for synthetic biology in the green alga *Chlamydomonas*. ACS Synthetic Biology 9: 1406–1417

Mentch SJ, Locasale JW (2016) One-carbon metabolism and epigenetics: understanding the specificity Annal NY Acad Sci 1363: 91–8

Mitchell AL, Attwood TK, Babbitt PC, Blum M, Bork P, Bridge A, Brown SD, Chang H-Y, El-Gebali S, Fraser MI et al. (2019) InterPro in 2019: improving coverage, classification and access to protein sequence annotations. Nucl Acids Res 47: D351–D360

Neupert J, Karcher D, Bock R (2009) Generation of *Chlamydomonas* strains that efficiently express nuclear transgenes. The Plant Journal 57: 1140–1150

Nielsen MJ, Rasmussen MR, Andersen CBF, Nexø E, Moestrup SK (2012) Vitamin B_12_ transport from food to the body’s cells—a sophisticated, multistep pathway. Nature Reviews Gastroenterology & Hepatology 9: 345–354

Ohwada K (1973) Seasonal cycles of vitamin B12, thiamine and biotin in Lake Sagami. Patterns of their distribution and ecological significance. Internationale Revue der gesamten Hydrobiologie und Hydrographie 58: 851–871

Orłowska M, Steczkiewicz K, Muszewska A (2021) Utilization of cobalamin is ubiquitous in early-branching fungal phyla. Genome Biology and Evolution 13: evab043

Panzeca C, Beck AJ, Tovar-Sanchez A, Segovia-Zavala J, Taylor GT, Gobler CJ, Sañudo-Wilhelmy SA (2009) Distributions of dissolved vitamin B_12_ and Co in coastal and open-ocean environments. Estuarine, Coastal and Shelf Science 85: 223–230

Pintner IJ, Altmeyer VL (1979) Vitamin B_12_-binder and other algal inhibitors Journal of Phycology 15: 391–398

Richter DJ, Berney C, Strassert JFH, Poh Y-P, Herman EK, Muñoz-Gómez SA, Wideman JG, Burki F, Vargas C de (2022) EukProt: A database of genome-scale predicted proteins across the diversity of eukaryotes. Peer Community Journal 2: e56

Ritz C, Baty F, Streibig JC, Gerhard D (2015) Dose-response analysis using R. PLoS One 10: e0146021

Rutsch F, Gailus S, Miousse IR, Suormala T, Sagné C, Toliat MR, Nürnberg G, Wittkampf T, Buers I, Sharifi A, et al (2009) Identification of a putative lysosomal cobalamin exporter altered in the cblF defect of vitamin B_12_ metabolism. Nature Genetics 41: 234–239

Sahni MK, Spanos S, Wahrman MZ, Sharma GM (2001) Marine corrinoid-binding proteins for the direct determination of vitamin B_12_ by radioassay. Analytical Biochemistry 289: 68–76

Sañudo-Wilhelmy SA, Gómez-Consarnau L, Suffridge C, Webb EA (2014) The role of B vitamins in marine biogeochemistry. Annual Review of Marine Science 6: 339–367

Shelton AN, Seth EC, Mok KC, Han AW, Jackson SN, Haft DR, Taga ME (2019) Uneven distribution of cobamide biosynthesis and dependence in bacteria predicted by comparative genomics. The ISME Journal 13: 789–804

Tang YZ, Koch F, Gobler CJ (2010) Most harmful algal bloom species are vitamin B_1_ and B_12_ auxotrophs. Proc Natl Acad Sci U S A 107: 20756–61

Warren MJ, Raux E, Schubert HL, Escalante-Semerena JC (2002) The biosynthesis of adenosylcobalamin (vitamin B_12_). Natural product reports 19: 390–412

Xie B, Bishop S, Stessman D, Wright D, Spalding MH, Halverson LJ (2013) *Chlamydomonas reinhardtii* thermal tolerance enhancement mediated by a mutualistic interaction with vitamin B_12_-producing bacteria. ISME J 7: 1544–55

Yu Z, Geisler K, Leontidou T, Young REB, Vonlanthen SE, Purton S, Abell C, Smith AG (2021) Droplet-based microfluidic screening and sorting of microalgal populations for strain engineering applications. Algal Res 56: 102293

