## Supplementary material for "The conserved protein CBA1 is required for vitamin B_12_ uptake in different algal lineages": Sayer et al. Supplementary material

**Supplementary Materials**

**Methods**

**Design of constructs for CRISPR/Cas9 editing of P. tricornutum**

Plasmid pMLP2127 (Table S2), featuring Cas9-YFP, two sgRNAs and a zeocin resistance cassette was constructed as follows. A homology template was designed that contained 2 homology regions of approximately 800 bp from regions flanking the *P. tricornutum* *CBA1* (*PtCBA1)* ORF (Figure 1a), which were amplified from the genome and placed either side of a nourseothricin resistance cassette in plasmid pMLP2117. The level 1 plasmid encoding the Cas9-YFP expression cassette (pICH47742:PtFCP:Cas9YFP), the level 0 plasmid containing the PtU6 promoter used to drive expression of the sgRNAs (pCR8/GW:PtU6) and the plasmid used as a template to amplify the sgRNA scaffold (pICH86966::AtU6p::sgRNA_PDS) were gifts from Dr. Amanda Hopes and Prof Thomas Mock (University of East Anglia) and are available on Addgene. All primer sequences used in the cloning process can be found in Table S2.

**Algal dose-response assay**

The algal dose response assay used to screen and quantitatively assess UVM4::pAS_R1 transformant lines was performed in the following manner. Firstly, three cultures of each line were grown in 96-well microtiter plates containing 200 µl TAP media supplemented with different concentrations of paromomycin and vitamin B_12_ (Figure 2b); after four days, optical density was measured at 730 nm (OD730) using a FluoStar OPTIMA (BMG labtech) plate reader; and finally, the data generated was modelled using a 4 parameter logistic equation (Ritz et al., 2015).

**B_12_-BOIDIPY Imaging**

*C. reinhardtii* strains were incubated with 1 μM B_12_-BODIPY for 1 hour at room temperature. Cells were pelleted by centrifugation at 5000 g and washed with TAP media 3 times. *C. reinhardtii* strains were imaged in a confocal laser scanning microscope (TCS SP8, Leica Microsystems, Germany) with an HC PL APO CS2 40x/1.30 aperture oil-immersion lens. Images were taken using the sequential mode provided by the Leica LAS software, with the channel used for chlorophyll and brightfield detection being taken first and the channel used for B_12_-BODIPY detection taken second. The first image was acquired with excitation from a white light source at 476 nm at 6% power and emissions were detected between 674 - 688 nm; chlorophyll settings included 10% gain. Brightfield imaging used 699.1% gain and a 2.89% offset. Frames were captured with a line average of 16 and a frame accumulation of 1. The second image was acquired with excitation from a white light source at 589 nm at 2% power and emissions were detected between 607 - 620 nm with 500% gain. Frames were captured with a line average of 6 and a frame accumulation of 4. The overlay images were produced automatically by the Leica LAS software. Inkscape was used to increase the lightness and contrast of all the images in the same manner.

**Identification of TE insert**

Standard BLAST searches against the C. reinhardtii reference genome CC-503 v5 were performed using the BLASTN Tool integrated into the Phytozome database, the Plant Comparative Genomics portal of the Department of Energy's Joint Genome Institute (https://phytozome.jgi.doe.gov/pz/portal.html#). As comparison matrix BLOSUM62 was used with a default word length of 11 bp. In addition, gaps were allowed and filter query settings were set on. Multiple sequence alignments (MSA) were generated employing the progressive aligner MUSCLE (Edgar, 2004) which featured rapid sequence distance estimation using k-mer counting, implemented as part of the UGENE bioinformatic suite (v38.1, Okonechnikov et al., 2012). Default parameters, optimized for best accuracy were used. Consensus sequences from nucleotide or peptide alignments were extracted using UGENE’s default consensus mode. BLAST searches against the NCBI nucleotide collection and non-redundant protein archive were performed using the MEGABLAST and BLASTP tools provided via the NIH website (https://blast.ncbi.nlm.nih.gov/Blast.cgi) using default paraments. Analysis of protein sequences for Pfam matches were conducted using the sequence search function from the Pfam server (http://pfam.xfam.org) applying default settings. The Repbase TE library for *C. reinhardtii* and relatives (https://www.girinst.org/repbase/) as well as the manually curated repeat library generated by Craig et al. (2021) (repeat_lib_v3_2.volvocales) were used to annotated the identified repetitive element.

**Table S1** - **Table of strains used in this work**

Where no reference is listed the strain was obtained from the Culture Collection of Algae and Protozoa (CCAP), Oban, Scotland

| **Species** | **Type of Growth Media** | **Reference** | **Conditions** |
| --- | --- | --- | --- |
| *Phaeodactylum tricornutum* CCAP 1055/1 | F/2 media (Guillard, 1975) |  | 18ºC, light intensity of 50 µE·m^-2^·s^-1^ under a 16 h light / 8 h dark cycle and constant shaking at 110 rpm |
| *Phaeodactylum tricornutum* CCAP 1055/1 ∆CBA1-1; ∆CBA1-2; ∆CBA1-3; | F/2 media (Guillard, 1975) with nourseothricin or zeocin |  | 18ºC, light intensity of 50 µE·m^-2^·s^-1^ under a 16 h light / 8 h dark cycle and constant shaking at 110 rpm |
| *Salmonella typhimurium* AR3612 *cysG metE* | M9 Minimal medium with methionine (50 mg/L) and cysteine (50 mg/L) | Raux et al., 1996 | 37°C, shaking speed 220 rpm |
| *Chlamydomonas reinhardtii* UVM4 | Tris-acetate phosphate (TAP) medium with Kropat’s trace elements (Kropat et al., 2011) but excluding selenium | Neupert et al., 2009 | 25°C under a continuous or 16 h / 8 h light / dark cycle (shaking speed of 120 rpm, light intensity of 90 µmol.m^-2^.s^-1^) |
| *Chlamydomonas reinhardtii* UVM4::pAS_R1 (UVM4-T12) | Tris-acetate phosphate (TAP) medium with Kropat’s trace elements (Kropat et al., 2011) but excluding selenium. Paromomycin 5-50 µg/ml. | This work | 25°C under a continuous or 16 h / 8 h light / dark cycle (shaking speed of 120 rpm, light intensity of 90 µmol.m^-2^.s^-1^) |
| *Chlamydomonas reinhardtii* UVM4-T12::pHyg3 #IM1-7 | Tris-acetate phosphate (TAP) medium with Kropat’s trace elements (Kropat et al., 2011) but excluding selenium. Paromomycin 5-20 µg/ml, hygromycin 10-20 µg/ml. | This work | 25°C under a continuous or 16 h / 8 h light / dark cycle (shaking speed of 120 rpm, light intensity of 90 µmol.m^-2^.s^-1^) |
| *Chlamydomonas reinhardtii* CC-5325 cw15 mt- (cw15) | Tris-acetate phosphate (TAP) medium with Kropat’s trace elements (Kropat et al., 2011) but excluding selenium. | Li et al., 2016 | 25°C under a continuous or 16 h / 8 h light / dark cycle (shaking speed of 120 rpm, light intensity of 90 µmol.m^-2^.s^-1^) |
| *Chlamydomonas reinhardtii* LMJ-119922 and LMJ-042227 | Tris-acetate phosphate (TAP) medium with Kropat’s trace elements (Kropat et al., 2011) but excluding selenium. Paromomycin 5-20 µg/ml | Li et al., 2016 | 25°C under a continuous or 16 h / 8 h light / dark cycle (shaking speed of 120 rpm, light intensity of 90 µmol.m^-2^.s^-1^) |
| IM4::pAS_C2 | Tris-acetate phosphate (TAP) medium with Kropat’s trace elements (Kropat et al., 2011) but excluding selenium. Paromomycin 5-20 µg/ml, hygromycin 10-20 µg/ml, Spectinomycin 75 µg/ml. | This work | 25°C under a continuous or 16 h / 8 h light / dark cycle (shaking speed of 120 rpm, light intensity of 90 µmol.m^-2^.s^-1^) |
| *Chlamydomonas reinhardtii* LMJ-040682 and LMJ-135929 | Tris-acetate phosphate (TAP) medium with Kropat’s trace elements (Kropat et al., 2011) but excluding selenium. Paromomycin 5-20 µg/ml | Li et al., 2016 | 25°C under a continuous or 16 h / 8 h light / dark cycle (shaking speed of 120 rpm, light intensity of 90 µmol.m^-2^.s^-1^) |
| LMJ-040682::pAS_C2 | Tris-acetate phosphate (TAP) medium with Kropat’s trace elements (Kropat et al., 2011) but excluding selenium. Paromomycin 5-20 µg/ml, Spectinomycin 75 µg/ml, +/- 10 µM thiamine. | This work | 25°C under a continuous or 16 h / 8 h light / dark cycle (shaking speed of 120 rpm, light intensity of 90 µmol.m^-2^.s^-1^) |
| LMJ-040682::pAS_C3 | Tris-acetate phosphate (TAP) medium with Kropat’s trace elements (Kropat et al., 2011) but excluding selenium. Paromomycin 5-20 µg/ml, Spectinomycin 75 µg/ml +/- 10 µM thiamine. | This work | 25°C under a continuous or 16 h / 8 h light / dark cycle (shaking speed of 120 rpm, light intensity of 90 µmol.m^-2^.s^-1^) |
| WT12-metE7 mutant | Tris-acetate phosphate (TAP) medium with Kropat’s trace elements (Kropat et al., 2011) but excluding selenium | Helliwell et al 2015 | 25°C under a continuous or 16 h / 8 h light / dark cycle (shaking speed of 120 rpm, light intensity of 90 µmol.m^-2^.s^-1^) |

**Table S2 - Oligonucleotides and plasmids used in this study**

F and R refer to forward and reverse primers respectively.

| **Type** | **Name** | **Purpose** | **Sequence (if oligonucleotide)** | **Notes** |
| --- | --- | --- | --- | --- |
| Oligonucleotide | sgCBA1.1.fwd | *P. tricornutum* cloning primer | AAGGTCTCACGAGGACCTACCTCCTCTACCAGTGGTTTTAGAGCTAGAAATAGCAAG |  |
| Oligonucleotide | sgCBA1.2.fwd | *P. tricornutum* cloning primer | AAGGTCTCACGAGGCGTTTGCGAGAGACCCACGTGTTTTAGAGCTAGAAATAGCAAG |  |
| Oligonucleotide | sgRNA.rv | *P. tricornutum* cloning primer | TGGTCTCAAGCGTAATGCCAACTTTGTACAAG |  |
| Oligonucleotide | CBA1.5HR.fwd | *P. tricornutum* cloning primer | AAGGTCTCAGGAGTCGACCACCACTTATTCC |  |
| Oligonucleotide | CBA1.5HR.rv | *P. tricornutum* cloning primer | AAGGTCTCAAGCGTTCGGAGTATCCTGATGG |  |
| Oligonucleotide | CBA1.3HR.fwd | *P. tricornutum* cloning primer | AAGGTCTCAGGAGGACTACTCAGTCTTATACACGTATTG |  |
| Oligonucleotide | CBA1.3HR.rv | *P. tricornutum* cloning primer | AAGGTCTCAAGCGACACAGAAATGATGCCTC |  |
| Oligonucleotide | gCBA1.fwd | *P. tricornutum* genotyping primer | GTTTCCCCCAAGCCTTTG | White triangle in Fig.1 |
| Oligonucleotide | gCBA1.rv | *P. tricornutum* genotyping primer | CAGCAAGGACGCTATTCAGG | White triangle in Fig.1 |
| Oligonucleotide | gCBA1in.fwd | *P. tricornutum* genotyping primer | CGCTCTTCTCCCAAGGATG | Grey arrow in Fig.1 |
| Oligonucleotide | gCBA1in.rv | *P. tricornutum* genotyping primer | GATCTCGTCCAAGAAGCAAGG | Grey arrow in Fig.1 |
| Oligonucleotide | NAT.rv | *P. tricornutum* genotyping primer | AGTGAACACGACGCTGAAGG | Grey arrow with circle in Fig.1 |
| Oligonucleotide | METE F | RT-qPCR METE | CCGCTACAGCCAGACTTCA |  |
| Oligonucleotide | METE R | RT-qPCR METE | GTGACAGCGACACGAACGT |  |
| Oligonucleotide | ON_56 | RT-qPCR RACK1 | CGTCTGTGGGACCTGAACAC |  |
| Oligonucleotide | ON_57 | RT-qPCR RACK1 | GCTCGCCAATGGTGTACTTG |  |
| Oligonucleotide | ON_160 | LMJ_135929 knockout confirmation | TTGAAGACATTATCCTACTACAGCACCTTCAAGGTGA AAA TTTCAGAA TGCC |  |
| Oligonucleotide | ON_113 | LMJ_135929 knockout confirmation | TTGAAGACATAGGGGTGACGTACGCCACGCG |  |
| Oligonucleotide | ON_114 | LMJ-040682 knockout confirmation | TTGAAGACATCCCTTACGCCGTGGAGCCTTGC |  |
| Oligonucleotide | ON_116 | LMJ-040682 knockout confirmation | TTGAAGACA TGGTGCGCAGCACCGGCAG |  |
| Oligonucleotide | ON_135 | LMJ-119922 knockout confirmation | CACCACCAGCACCCAGTG |  |
| Oligonucleotide | ON_141 | LMJ-119922 knockout confirmation | GCTCCCGAACCCGTCAC |  |
| Oligonucleotide | ON_177 | LMJ-042227 knockout confirmation | AGCAGCAGTAGAAGCAGCG |  |
| Oligonucleotide | ON_178 | LMJ-042227 knockout confirmation | GGCTGTACTCCGCCTCCAT |  |
| Oligonucleotide | ON_121 | *Golden gate Cre02.g081050* promoter | TTGAAGACATGGAGCAGCAGTCGCTGTGTCTCGTACTCCAC | Into G2 backbone vector |
| Oligonucleotide | ON_151 | *Golden gate Cre02.g081050* promoter | TTGAAGACATACACATTTGACACAATGTCGTTTCCCAAGTTTCAGGG | Into G2 backbone vector |
| Oligonucleotide | ON_152 | *Golden gate Cre02.g081050* promoter | TTGAAGACATGTGTTCACGCCATGCGCC | Into G2 backbone vector |
| Oligonucleotide | ON_153 | *Golden gate Cre02.g081050* promoter | TTGAAGACATGGTGTCGGGGGAGGCCTTG | Into G2 backbone vector |
| Oligonucleotide | ON_154 | *Golden gate Cre02.g081050* promoter | TTGAAGACATCACCTCGTTCTCCTTGTCAGCCACGCAATCGCAAGGTTGGATCTTC | Into G2 backbone vector |
| Oligonucleotide | ON_155 | *Golden gate Cre02.g081050* promoter | TTGAAGACATGGTATCGCCACTACCGCAACCCAGTCCGC | Into G2 backbone vector |
| Oligonucleotide | ON_156 | *Golden gate Cre02.g081050* promoter | TTGAAGACATTACCCCGTGCTCCTCGC | Into G2 backbone vector |
| Oligonucleotide | ON_122 | *Golden gate Cre02.g081050* promoter | TTGAAGACATCATTCACAATGTATGTGTAGCGCAACCTTGC | Into G2 backbone vector |
| Oligonucleotide | ON_123 | *Golden gate Cre02.g081050* terminator | TTGAAGACATGCTTGCGCCCCGCCTCCCAG | Into D2 backbone vector |
| Oligonucleotide | ON_161 | *Golden gate Cre02.g081050* terminator | TTGAAGACATAGATCGCGGGACCACCGTCTACC | Into D2 backbone vector |
| Oligonucleotide | ON_162 | *Golden gate Cre02.g081050* terminator | TTGAAGACATATCTCATGACTAACTGAAAGGTGCGGCGTGG | Into D2 backbone vector |
| Oligonucleotide | ON_124 | *Golden gate Cre02.g081050* terminator | TTGAAGACATAGCGAGCAATGTTCTATTGTTTGCGGGGGATTCGG | Into D2 backbone vector |
| Oligonucleotide | ON_112 | *Golden gate Cre02.g081050* CDS-S | TTGAAGACATAATGGCGTCTCGCGGCCTCTT | Amplified from Cre02.g081050 CDS-NS template to ensure no BpiI or BsaI sites into H2 backbone vector |
| Oligonucleotide | ON_119 | *Golden gate Cre02.g081050* CDS-S | TTGAAGACATAAGCTTACAGGCCCCAGCCCAGCACG | Amplified from Cre02.g081050 CDS-NS template to ensure no BpiI or BsaI sites into H2 backbone vector |
| Oligonucleotide | ON_112 | *Golden gate Cre02.g018050* CDS-NS | TTGAAGACATAATGGCGTCTCGCGGCCTCTT | Into E1 backbone vector |
| Oligonucleotide | ON_157 | *Golden gate Cre02.g018050* CDS-NS | TTGAAGACATACACATTCACGCCGGTGACCTTCAG | Into E1 backbone vector |
| Oligonucleotide | ON_158 | *Golden gate Cre02.g018050* CDS-NS | TTGAAGACATGTGTTCCCCACGACGCG | Into E1 backbone vector |
| Oligonucleotide | ON_159 | *Golden gate Cre02.g018050* CDS-NS | TTGAAGACATGATACCTCGAAGTTCTGAGCCAC | Into E1 backbone vector |
| Oligonucleotide | ON_160 | *Golden gate Cre02.g018050* CDS-NS | TTGAAGACATTATCCTACTACAGCACCTTCAAGGTGAAAATTTCAGAATGCC | Into E1 backbone vector |
| Oligonucleotide | ON_120 | *Golden gate Cre02.g018050* CDS-NS | TTGAAGACATACCTCCCAGGCCCCAGCCCAGCACG | Into E1 backbone vector |
| Oligonucleotide | ON_104 | pHyg3 insertion confirmation | CCATCGCTGTCACTGGGT | *Cre12.g508644* across insertion |
| Oligonucleotide | ON_106 | pHyg3 insertion confirmation | GATGACGCAGACTTTGCCAC | *Cre12.g508644* across insertion |
| Oligonucleotide | ON_104 | pHyg3 insertion confirmation | CCATCGCTGTCACTGGGT | *Cre12.g508644* left junction |
| Oligonucleotide | ON_89 | pHyg3 insertion confirmation | TACGGTCGAGAAGTAACAGGGA | *Cre12.g508644* left junction |
| Oligonucleotide | ON_79 | pHyg3 insertion confirmation | AAAACCCTGGCGTTACCCAA | *Cre12.g508644* right junction |
| Oligonucleotide | ON_106 | pHyg3 insertion confirmation | GATGACGCAGACTTTGCCAC | *Cre12.g508644* right junction |
| Oligonucleotide | ON_205 | Cre02.g081050 disruption confirmation | CCTTGTCAGCCACGCAATC | *Cre02.g081050* left junction |
| Oligonucleotide | ON_201 | Cre02.g081050 disruption confirmation | GCTTCCGGGCGAGAACCTTATG | *Cre02.g081050* left junction |
| Oligonucleotide | ON_208 | Cre02.g081050 disruption confirmation | GCAAGCAGACGGAGGACAAC | *Cre02.g081050* right junction |
| Oligonucleotide | ON_203 | Cre02.g081050 disruption confirmation | CGTGTGCAAAGGGCCATG | *Cre02.g081050* right junction |
| Plasmid | pMLP2127 | *P. tricornutum* CRISPR-Cas9 |  | Level 2 CRISPR-Cas9 construct |
| Plasmid | pICH47742:PtFCP:Cas9YFP | *P. tricornutum* CRISPR-Cas9 |  | Level 1 plasmid – Cas9-YFP |
| Plasmid | pCR8/GW:PtU6 | *P. tricornutum* CRISPR-Cas9 |  | Level 0 - PtU6 promoter |
| Plasmid | pICH86966::AtU6p::sgRNA_PDS | *P. tricornutum* CRISPR-Cas9 |  | Amplify the sgRNA scaffold |
| Plasmid | pMLP2117 | *P. tricornutum* CRISPR-Cas9 |  | Homology template |
| Plasmid | pAS_R1 | B_12_ reporter |  | *P_METE_-AphVIII-T_CA1_* |
| Plasmid | pAS_C2 | Complementation |  | *P_PSAD_-aada-T_PSAD_ P_Cre02.g081050_-Cre02.g081050-mVenus-T_Cre02.g081050_* |
| Plasmid | pAS_C3 | Complementation |  | *P_PSAD_-aada-T_PSAD_ P_Rbcs2/Hsp70_-CrTHI4-4N-Cre02.g081050-T_CA1_* |
| Plasmid | pHyg3 | Insertional cassette |  | *P_beta2-tubulin_-AphVII-T_Rbcs2_* |

**Supplementary references**

**Capella-Gutiérrez S, Silla-Martı́nez JoséM, Gabaldón T** (2009) [trimAl: A tool for automated alignment trimming in large-scale phylogenetic analyses](https://doi.org/10.1093/bioinformatics/btp348). Bioinformatics **25**: 1972–1973

**Craig RJ, Hasan AR, Ness RW, Keightley PD** (2021) [Comparative genomics of *Chlamydomonas*](https://doi.org/10.1093/plcell/koab026). Plant Cell **33**:1016-1041

**Edgar RC** (2004) [MUSCLE: multiple sequence alignment with high accuracy and high throughput](https://academic.oup.com/nar/article/32/5/1792/2380623?login=true). Nucleic Acids Res 32, 1792–1797 (2004).

**Guillard RRL** (1975) [Culture of phytoplankton for feeding marine invertebrates](https://doi.org/10.1007/978-1-4615-8714-9%7b\_%7d3). *In* WL Smith, MH Chanley, eds, Culture of marine invertebrate animals: Proceedings —1st conference on culture of marine invertebrate animals greenport. Springer US, Boston, MA, pp 29–60

**Katoh K, Standley DM** (2013) [MAFFT multiple sequence alignment software version 7: Improvements in performance and usability](http://mbe.oxfordjournals.org/content/30/4/772.abstract). Molecular Biology and Evolution **30**: 772–780

**Kelley LA, Mezulis S, Yates CM, Wass MN, Sternberg MJE** (2015) [The Phyre2 web portal for protein modeling, prediction and analysis](https://doi.org/10.1038/nprot.2015.053). Nat Protoc **10**: 845–58

**Letunic I, Bork P** (2019) [Interactive tree of life (iTOL) v4: Recent updates and new developments](https://doi.org/10.1093/nar/gkz239). Nucleic Acids Research **47**: W256–W259

**Nguyen L-T, Schmidt HA, Haeseler A von, Minh BQ** (2015) [IQ-TREE: A fast and effective stochastic algorithm for estimating maximum-likelihood phylogenies](https://doi.org/10.1093/molbev/msu300). Molecular Biology and Evolution **32**: 268–274

**Okonechnikov K, Golosova O, Fursov M, Team U** (2012) [Unipro UGENE: a unified bioinformatics toolkit](https://academic.oup.com/bioinformatics/article/28/8/1166/195474). Bioinformatics 28, 1166–1167

**Raux-Deery E, Lanois A, Levillayer F, Warren MJ, Brody E, Rambach A, Thermes C** (1996) [*Salmonella typhimurium* cobalamin (vitamin B_12_) biosynthetic genes: Functional studies in *S. typhimurium* and *Escherichia coli.*](https://doi.org/10.1128/jb.178.3.753-767.1996) Journal of Bacteriology **178**: 753–67

**Ritz C, Baty F, Streibig JC, Gerhard D** (2015) [Dose-response analysis using R](https://doi.org/10.1371/journal.pone.0146021). PLoS One **10**: e0146021

**Figure S1.** **Characterisation of B_12_ uptake in *C. reinhardtii* insertional mutant lines.** To determine whether any of the 7 insertional mutant lines (IM) isolated from mutagenesis of UVM4::T12 showed impaired B_12_ uptake, the B_12_-uptake assay was performed as described in the materials and methods. Control-1, Control-2 and Control-3 lines were picked from solid media with paromomycin and hygromycin but without vitamin B_12_, whereas IM1-IM7 lines were picked from solid media containing paromomycin, hygromycin and vitamin B_12_. The total was inferred by the addition of the cell and media fractions. The dashed line indicates the amount of B_12_ added in the uptake assay. Standard deviation error bars are shown, n=4. Statistical analysis was performed on the media fraction, and Tukey’s test identified the following comparisons to be significantly different from one another (only reporting IM strains different from UVM4, UVM4-T12 or Control-[1,2,3] strains): IM4 vs UVM4 (p<1e^-12^), UVM4-T12 (p<1e^-12^), Control-1 (p<1e^-12^), Control-2 (p<1e^-12^), Control-3 (p<1e^-12^); and IM2 vs UVM4 (p<0.05).

**Figure S2. Visualisation of B_12_-BODIPY uptake in *C. reinhardtii* using confocal microscopy.** To assess B_12_ uptake more directly, the insertional mutant line IM4 (impaired B_12_ uptake) and its parental line UVM4-T12 were incubated with the fluorescent B_12_ analogue B_12_-BODIPY and the samples were imaged using confocal microscopy, as described in the materials and methods. Channels shown are brightfield (greyscale), chlorophyll (red), B_12_-BODIPY (yellow) and an overlay.
